# Productivity of the eelgrass biotope and the resilience of fisher-hunter-gatherers in prehistoric Denmark

**DOI:** 10.1101/2024.05.06.592700

**Authors:** John Meadows, Anders Fischer

## Abstract

In the shallow coastal waters of the Danish archipelago, eelgrass (*Zostera marina*) forms extensive and extremely productive stands, which support food webs that are isotopically distinct from those based on the primary productivity of marine phytoplankton. The isotopic signatures of eelgrass and phytoplankton food webs explain much of the variation in δ^13^C, δ^15^N and δ^34^S values in marine vertebrate remains recovered from prehistoric coastal sites in Denmark. Isotope data from human and dog remains reflect the overwhelming importance of the eelgrass biotope to the fisher-hunter-gatherer (FHG) subsistence during the Late Mesolithic Ertebølle epoch (∼7400-5800 cal BP). By recognising eelgrass and pelagic resources as isotopically distinct, we obtain more accurate diet reconstructions and dietary ^14^C reservoir effect corrections for Ertebølle individuals. Longstanding research issues, including technology, palaeodemography, and the resilience and sustainability of the FHG economy need to be reassessed in view of the isotopic signature and productivity of the eelgrass biotope in Danish waters. Eelgrass productivity is also pertinent to comparisons between Ertebølle and Neolithic economies in Denmark, and between FHG adaptations in Denmark and neighbouring regions, and may be key to understanding the much-discussed delay in the transition to farming.

## 1. Introduction

### 1.1. Isotopes and the Ertebølle economy

The numerous shell middens and other coastal settlements in Denmark play a fundamental role in understanding fisher-hunter-gatherer (FHG) lifeways in Mesolithic Europe. Already in the 1800s, researchers associated the Late Mesolithic Ertebølle epoch with intensive exploitation of coastal resources (Fischer and Kristensen 2002). In his ground-breaking study of dietary stable isotopes, Henrik Tauber (1981) showed a sharp contrast in collagen δ^13^C between Ertebølle and Neolithic human bones from coastal settings. There were no C_4_ plants in this region before the Bronze Age introduction of millet, so higher δ^13^C values indicated regular consumption of marine resources; Ertebølle δ^13^C values suggested diets like those of Greenland Inuit, while Neolithic farmers apparently relied mainly or entirely on terrestrial resources. Tauber (1984) noted the implications for dating Ertebølle skeletons, namely that their conventional (fractionation-corrected) radiocarbon (^14^C) ages could be too old by up to ∼400 years, the apparent ^14^C age of carbonate in surface ocean water. Tauber realised that δ^13^C could be used to approximate the proportion of ^14^C in human bones derived from marine resources, and thus to estimate individual dietary reservoir effects (DREs), but cautioned that the wide δ^13^C range in marine species meant large uncertainties in DRE corrections.

Subsequent discussion focussed on how complete, rapid and universal was the shift from marine to terrestrial diets at the Mesolithic-Neolithic transition (Milner et al. 2004; Richards et al. 2003), and on whether Ertebølle people moved between coastal and inland areas (Fischer et al. 2007a; Noe-Nygaard 1995). Tauber’s studies drew attention to the under-representation of fish in older excavations, and research evolved to address this issue. Recovery and analysis of fish remains was applied systematically (Enghoff 2011; Gron and Robson 2016; Ritchie 2010), and there was increasing interest in wooden fishing structures, often preserved in marine sediments next to Ertebølle culture (EBC) coastal settlements, which would have enabled harvesting of fish stocks from large expanses of shallow water (Fischer 2007; Pedersen 1995; Pedersen et al. 2013). As early as 1994, Inge Bødker Enghoff realised that almost all the fish found at EBC coastal sites might have been caught in eelgrass meadows: ‘Some of the species represented spend their entire life close to the coast, e.g. in the eel-grass zone…The garfish, for example, breeds in the eel-grass zone during summer … Even the spurdog enters shallow water, including the eel-grass zone’ (Enghoff 1994).

We now see that garfish (*Belone belone*) and probably spurdog (*Squalus acanthias*) were isotopically offshore (‘pelagic’) species, regardless of where they were caught (see 3.2.1), but in 1994, there were almost no stable isotope data for prehistoric fish and marine mammals from Denmark. As these data appeared, the wide range of δ^13^C values was attributed either to movement between the brackish Baltic and saline North Sea (Craig et al. 2006), or to fishing in estuarine locations, or of catadromous species (Robson et al. 2012). These factors will have contributed to δ^13^C variation, but should only give rise to lower (more negative) δ^13^C than in fully marine species, yet Danish fish are notable for their elevated δ^13^C values (3.2.1).

We argue that δ^13^C variation in Danish marine fauna is due primarily to the productivity of the eelgrass (*Zostera marina*) biotope in this region. Archaeologists have paid little attention to the long history of eelgrass research in the Danish archipelago by marine biologists (Krause-Jensen et al. 2021). Only Pleuger and Makarewicz (2020), in their analysis of remains from Kainsbakke and Kirial Bro, two Pitted Ware (Middle Neolithic, ∼5000 cal BP) sites in the Djursland peninsula (Fig. 1), proposed a link between δ^13^C values of fish and *Zostera*. Had they also reviewed evidence from EBC sites, they might have attached more significance to this association.

**Figure 1:**
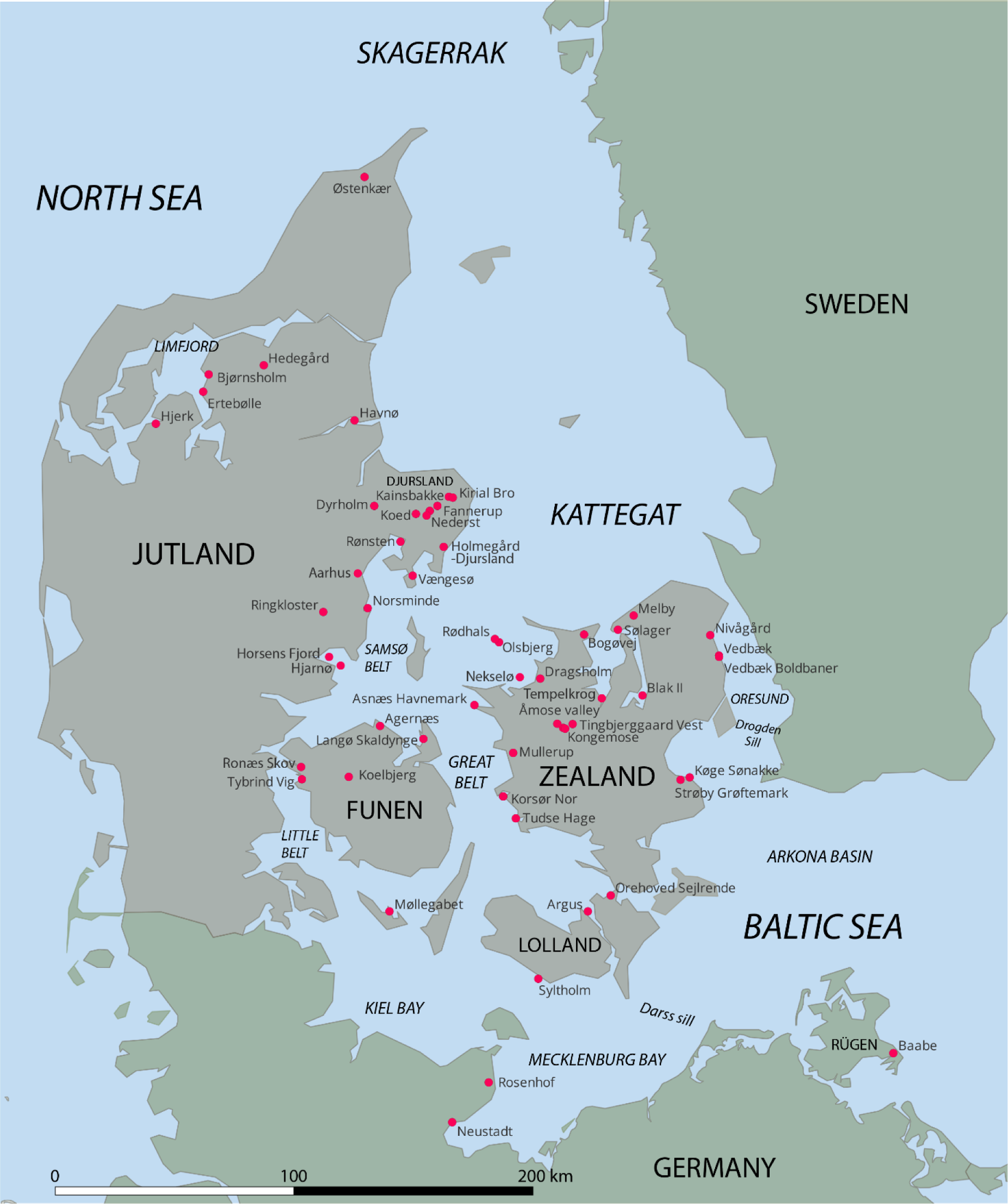
Modern coastline of Denmark and neighbouring areas of Germany and Sweden. Created in QGIS 3.28. Geographic coordinates for archaeological sites shown are provided in Supplementary Tables 1-3.

We found a similar isotopic pattern to Pleuger and Makarewicz (2020) in Ertebølle fauna at Rødhals (Fischer et al. 2021, appendix B). Now, we argue that, given patterns in domestic dog and human stable isotopes (3.2.2, 3.2.3), the Ertebølle economy in much of Denmark was based on the productivity and predictability of marine resources from the eelgrass biotope. The productivity of eelgrass therefore affects human diet reconstruction and DRE correction models (3.3.2, 3.3.3), and must be considered in explaining the success, resilience and eventual demise of the Ertebølle system. It may also be relevant to subsequent adaptations in Denmark, and to the trajectories of FHG economies elsewhere in the Baltic region (3.4).

### 1.2. The modern eelgrass biotope

Seagrasses, and particularly eelgrass, form dense stands on the seabed in shallow (<20m depth) coastal waters in many parts of the globe, providing sheltered habitats for a wide range of invertebrates and young fish. Seagrass meadows are key resource areas for traditional low-technology fishing communities (Unsworth et al. 2019), and ethnoarchaeological and archaeobotanical research suggests that eelgrass rhizomes could have been consumed by coastal indigenous groups in North America (Cullis-Suzuki 2007; Fauvelle et al. 2012).

Danish coastal waters, including the southwest Baltic, Belt Sea and Kattegat, form a shallow sill between the Baltic proper and the Skagerrak (Fig. 1). More freshwater enters the Baltic from precipitation and river catchments than leaves it by evaporation, providing a large net outflow to the North Sea, which limits the salinity in Danish waters and creates ideal conditions for eelgrass to flourish. The eelgrass zone was traditionally a highly productive and reliable fishing ground (Johansson 1999; Petersen 1899), but eelgrass meadows in Danish waters now cover only 10-20% of their extent in the early 1900s (Boström et al. 2014). Disease caused a dramatic loss of eelgrass in the 1930s (Rasmussen 1977), with recovery limited by eutrophication.

#### 1.2.1. Eelgrass biotope productivity

Studying eelgrass at Hvidøre in the Oresund in 1978-79, Wium-Andersen and Borum (1984) reported that ‘annual eelgrass production … is at the same level as that of agricultural crops’, and nine times higher than phytoplankton productivity in the same area. Phytoplankton biomass was much greater overall, due to the limited area of eelgrass meadows in the Oresund, but eelgrass accounted for most of the primary production in shallow water. Other macrophytes occur in eelgrass meadows, but eelgrass shoot density is particularly high in Danish waters, and total biomass is therefore dominated by this species (Boström et al. 2014; Wium-Andersen and Borum 1984). Boström et al. (2014) found that eelgrass summer growth (daily increase in biomass dry weight per square metre) was far higher at sites in the Kattegat/Belt Sea (7-18g DW m^2^ d^-1^) than in the Skagerrak (2-5g DW m^2^ d^-1^) or Baltic proper (<1-2g DW m^2^ d^-1^). If total annual biomass formation is 170% of summer growth (estimate based on Sand-Jensen (1975)), eelgrass net primary productivity (NPP) in Danish waters would be ∼500-1200 gC m^2^ a^-1^ (excluding epiphytes). The Hvidøre site is typical (measured NPP 814 gC m^2^ a^-1^, or 884 gC m^2^ a^-1^ including epiphytes (Wium-Andersen and Borum 1984)). Danish eelgrass meadows are near the top of the NPP range in terrestrial biomes used by modern foragers (Zhu et al. 2021): of 357 recorded biomes, 233 have an NPP <500 gC m^2^ a^-1^ and only 29 have an NPP >1200 gC m^2^ a^-1^.

Krause-Jensen et al. (2003) found that light, wave exposure and salinity all limited the density and extent of natural eelgrass stands in Danish waters. Light was the most important factor. Anything reducing light penetration (wave action, sedimentation, phytoplankton blooms) affects the survival and productivity of eelgrass (Moore et al. 1997). Light penetration decreases with water depth, but wave action in shallow water increases turbidity and uproots plants, so maximum eelgrass biomass is at about 4m depth. On exposed coasts, continuous stands of eelgrass are rare at depths of less than ∼2m, but in protected fjords and inlets, they can occur in shallower water. The optimal water temperature for eelgrass growth is 10-20 °C (Nejrup and Pedersen 2008). Productivity may decline above 21 °C (the average August temperature today in shallow waters of the Kattegat and Belt Sea); water below 5 °C sharply reduces productivity but is not fatal, but above 25 °C mortality increases dramatically (Olesen and Sand-Jensen 1993).

Eelgrass productivity peaks in moderately saline water (Nejrup and Pedersen 2008). Zhang et al. (2022) found maximum *Zostera marina* survivorship at 18-21‰ salinity, almost exactly the modern range in internal Danish coastal waters, which may explain the exceptionally high eelgrass productivity in this region, which includes the southwest Baltic, west of the Darss Sill (Fig. 1). In the Baltic proper, from the Arkona basin eastwards, salinity is <10‰ (Lehmann et al. 2022), and eelgrass productivity is consequently much lower than in the Kattegat-southwest Baltic. Eelgrass productivity in the Skagerrak may be limited by a combination of higher salinity, more wave exposure and greater tidal range than in the Kattegat.

#### 1.2.2. Eelgrass isotope signature

Eelgrass leaves, stems and rhizomes have much higher (less negative) δ^13^C values than marine phytoplankton, due to differences in fractionation during photosynthesis of Dissolved Inorganic Carbon (DIC, δ^13^C ∼0‰). Fractionation from DIC to tissue (Δ^13^C) is ∼ -20‰ in phytoplankton (Descolas-Gros and Fontugne 1990), but Δ^13^C in eelgrass averages -9.1±2.2‰ (Papadimitriou et al. 2005). In part, this is due to rapid growth depleting the limited supply of CO_2-_, and the capacity of eelgrass to photosynthesise the more abundant and isotopically heavier HCO_3-_ (due to kinetic effects, isotopic equilibrium between HCO_3-_ and CO_2-_ in DIC is reached when CO_2-_ δ^13^C is ∼8‰ lower) (McPherson et al. 2015; Papadimitriou et al. 2005). Eelgrass δ^13^C is therefore even higher in summer (Kim et al. 2014; Tanaka et al. 2008), when most eelgrass biomass is formed, particularly at higher latitudes. Since the 1970s, δ^13^C of the organic fraction of marine sediment has been used to show what proportion of its carbon content was photosynthesised by eelgrass (Fry et al. 1977; Röhr et al. 2018), and ecologists have used δ^13^C in invertebrates, fish and seals to estimate the contributions of eelgrass and phytoplankton to marine food webs (Burton and Koch 1999; Fry et al. 1977; McConnaughey and McRoy 1979; Thayer et al. 1978).

Through a process driven by sulfate-reducing bacteria in the seabed (McGlathery et al. 1998), eelgrass and other seagrass rhizomes can fix nitrogen from the sediment, which is not available to phytoplankton, and which is typically depleted in ^15^N relative to oceanic NH^4+^ (Russell et al. 2018). This process could reinforce negative correlations between δ^13^C and δ^15^N observed in several fish taxa from the eelgrass biotope (3.2.1). Eelgrass also has a distinct sulfur-isotope (δ^34^S) signature. δ^34^S values in pelagic species should approach those in marine sulfate (+21‰; Rees et al. (1978)), but seagrasses absorb and metabolise ^34^S-depleted sulfide generated by sulfate-reducing bacteria in sediment (Fry et al. 1982). *Zostera* roots and rhizomes take up ^34^S-depleted sulfide, and their δ^13^C and δ^34^S values vary according to stand density (Frederiksen et al. 2006; Hasler-Sheetal and Holmer 2015; Holmer et al. 2009). δ^34^S in combination with δ^13^C have been used to quantify the contribution of *Zostera* to marine invertebrate diets (Kharlamenko et al. 2001).

#### 1.2.3. Eelgrass food webs

Eelgrass primary productivity is archaeologically important to the extent that eelgrass biomass was ultimately converted into potential food resources. Ideally, by tracking the isotopic signature of eelgrass through the modern food web, we could quantify its contribution to the sustainable offtake of species directly consumed by Mesolithic humans and dogs (shellfish, eels, young cod etc.). Modern eelgrass faunal communities in Danish waters are probably unrepresentative of the potential fauna (Boström et al. 2014), however, and modern studies often aim to assess the impact of eutrophication, which dramatically reduces eelgrass productivity (Olsen et al. 2011). A food-web isotope study of two sites on Funen (Thormar et al. 2016) showed that small fish species depended on crustaceans, whose diet was derived mainly from phytoplankton and macroalgae, but <10% of sediment organic carbon was eelgrass-derived. A similar situation applies to a study in Odense Fjord (Kristensen et al. 2019). Mittermayr et al. (2014) found that invertebrates in an eelgrass meadow in Kiel Bay relied mainly on ‘seston’ (fine particulate matter in suspension), or to a lesser extent on epiphytes, not eelgrass itself; epiphyte, seston and invertebrate δ^34^S values were similar, and much higher than eelgrass δ^34^S, incidentally confirming that eelgrass takes up sulfide from the sediment. The site was in an exposed location, however, and eelgrass detritus was regularly washed out. In all three cases, the main food for most crustaceans was organic detritus, which elsewhere is derived mainly from decaying eelgrass (Röhr et al. 2018). Isotopic evidence from modern invertebrates, fish and seals in the North Pacific (Burton and Koch 1999; Kharlamenko et al. 2001; McConnaughey and McRoy 1979) is inexplicable without substantial direct or indirect consumption of *Zostera* biomass.

## 2. Materials and methods

To test the hypothesis that detrital *Zostera* (leaves and below-ground biomass) was an important food source for marine invertebrates and indirectly for the species which consumed them, we sampled nine domestic dogs (*Canis familiaris*) and three otters (*Lutra lutra*) found at five EBC sites in Denmark (Appendix A), and obtained ^14^C ages and stable isotope values (δ^13^C, δ^15^N, δ^34^S) for comparison with published data. If eelgrass biomass was a major food source, we expected that its distinct isotopic signature (1.2.2) would lead to correlations between consumer δ^13^C, δ^15^N, and δ^34^S values at higher trophic levels. If elevated δ^13^C values in Mesolithic dogs and otters are associated with lower δ^15^N and particularly lower δ^34^S values, isotopic variation in coastal dogs should primarily reflect the roles of pelagic and eelgrass food webs, and show that eelgrass itself, not epiphytes, was the main food source within the eelgrass biotope.

All sampled bones are stored at the Zoological Museum in Copenhagen. Bone powder or crushed fragments of all bones were extracted at the Leibniz Laboratory, Christian-Albrechts University (Kiel, Germany) following Grootes et al. (2004), and part of the freeze-dried collagen obtained was combusted, graphitised and dated by Accelerator Mass Spectrometry (AMS), following Nadeau et al. (1998). In one case, the Østenkær dog, two graphite targets were measured, for routine quality control purposes. Part of the extracted collagen from each sample was combusted and analysed in quadruplicate by Elemental Analysis-Isotope Ratio Mass Spectrometry (EA-IRMS) at isolab GmbH (Schweitenkirchen, Germany), following Sieper et al. (2006).

We analysed new and published data using Past 4.10 (Hammer et al. 2001) for basic statistics, FRUITS 2 (Fernandes et al. 2014) for diet reconstruction, and OxCal 4.4 (Bronk Ramsey 2009) for ^14^C modelling. In Past 4.10, we performed bivariate regressions on isotope data (δ^13^C vs δ^15^N, δ^13^C vs δ^34^S, and δ^15^N vs δ^34^S) using Reduced Major Axis (RMA) regression, which, unlike ordinary least square regression, assumes that both variables contain measurement uncertainty, and minimises the sum of both vertical and horizontal distances of the standardised data points from the regression line. We used k-means clustering to segment the dog and human isotope data, in order to focus on cases in which the isotopic influence of terrestrial and freshwater biotopes was minimal. Three clusters were necessary to select dogs and humans with overwhelmingly marine diets, for comparison with fish and marine mammal isotope data.

## 3. Results and Discussion

### 3.1. New analytical data

Newly obtained results are given in Table 1. Collagen yields (3.1-18.2% of bone weight) are all good and atomic C/N, C/S and N/S ratios are within accepted ranges for well-preserved collagen (Guiry and Szpak 2021; Nehlich and Richards 2009). Isotopic results are not correlated with collagen yields or atomic ratios, and should therefore represent endogenous values rather than diagenesis (Guiry and Szpak 2021). The paired results from the Østenkær dog, KIA-57421, are statistically consistent with a mean ^14^C age of 5423±27 BP (T=0.9, T’(5%)=3.8, df=1) (Ward and Wilson 1978). The Østenkær dog was dated previously (LuS-6618, 5400±50 BP (Enghoff 2011)), as was one of the otter bones, from Rødhals (UCIAMS-230939, 5300±20 BP (Fischer et al. 2021)); both of our new dates are compatible with the published ^14^C ages. Given their ^14^C ages, all 12 bones are associated with the EBC.

**Table 1:** New analytical data from bone collagen. Species determinations by I.B. Enghoff (Østenkær) and A.B. Gotfredsen (all others). Each sample can be attributed to a different individual.

| Lab code | Sample identifier | site | taxon | yield (%) | %C | %N | %S | C/N | C/S | N/S | $\delta^{13}\text{C}$ | $\delta^{15}\text{N}$ | $\delta^{34}\text{S}$ | $^{14}\text{C}$ age |
| --- | --- | --- | --- | --- | --- | --- | --- | --- | --- | --- | --- | --- | --- | --- |
| KIA-57002 | P177 - 104/1968 | Olsbjerg | dog | 12.6 | 38.7 | 14.1 | 0.20 | 3.21 | 517 | 161 | -10.81 | 12.30 | 6.39 | 5815±40 |
| KIA-57003 | P178 - 104/1968 | Olsbjerg | dog | 6.4 | 38.6 | 14.1 | 0.24 | 3.20 | 429 | 134 | -8.47 | 11.15 | 5.54 | 5785±40 |
| KIA-57004 | P179 - 104/1968 | Olsbjerg | dog | 9.8 | 38.4 | 14.0 | 0.21 | 3.21 | 487 | 152 | -8.13 | 11.03 | 5.24 | 5680±35 |
| KIA-57005 | P180 - 104/1968 | Olsbjerg | dog | 10.9 | 38.0 | 14.0 | 0.22 | 3.17 | 460 | 145 | -6.74 | 10.33 | 1.47 | 5720±35 |
| KIA-57006 | P181 - 104/1968 | Olsbjerg | dog | 5.5 | 38.4 | 14.0 | 0.24 | 3.20 | 427 | 133 | -10.78 | 13.81 | 7.48 | 5870±40 |
| KIA-57012 | P187 - 5/2014 | Asnæs Havnemark | dog | 4.8 | 35.4 | 13.2 | 0.19 | 3.14 | 497 | 158 | -9.85 | 11.45 | 7.65 | 5670±40 |
| KIA-57013 | P188 - 5/2014 | Asnæs Havnemark | dog | 5.5 | 36.2 | 13.5 | 0.21 | 3.13 | 460 | 147 | -10.49 | 13.83 | 8.60 | 5855±40 |
| KIA-57010 | P185 - 5/2014 | Asnæs Havnemark | otter | 3.1 | 32.7 | 12.2 | 0.25 | 3.12 | 348 | 112 | -10.57 | 13.83 | 7.15 | 5525±35 |
| KIA-57011 | P186 - 5/2014 | Asnæs Havnemark | otter | 4.1 | 34.4 | 12.8 | 0.22 | 3.15 | 417 | 132 | -15.38 | 12.42 | 8.39 | 5550±40 |
| KIA-57018 | P435 - 17/1938 | Rødhals | otter | 18.2 | 38.2 | 14.2 | 0.28 | 3.14 | 364 | 116 | -7.70 | 11.57 | 6.23 | 5285±35 |
| KIA-57420 | ZMK 23/1982 SVM 2002022 x217 100/263 | Bodal K (Knoglebo) | dog | 4.3 | 40.1 | 14.8 | 0.28 | 3.17 | 382 | 120 | -20.05 | 8.46 | 6.86 | 5251±34 |
| KIA-57421 | x527 491/613 lag 8 | Østenkær | dog | 6.0 | 40.3 | 14.6 | 0.3 | 3.23 | 358 | 111 | -21.21 | 7.79 | 9.14 | 5448±37; 5398±37 |

### 3.2. Patterns in archaeological stable isotopes

#### 3.2.1. Marine taxa

There are published δ^13^C and δ^15^N values for multiple individuals of the marine vertebrate taxa most frequently found in Ertebølle cultural contexts in Denmark and the southwest Baltic, often from several sites (Craig et al. 2006; Fischer et al. 2007a; Pleuger and Makarewicz 2020; Robson et al. 2016). Figure 2 shows δ^13^C and δ^15^N values for marine fish bones from Danish prehistoric sites. Normally, δ^13^C and δ^15^N increase with trophic level, within food webs and even within species (Jennings et al. 2002), so that δ^13^C and δ^15^N values are positively correlated. Here, however, higher δ^15^N values are associated with lower δ^13^C values, overall and within most taxa.

**Figure 2.**
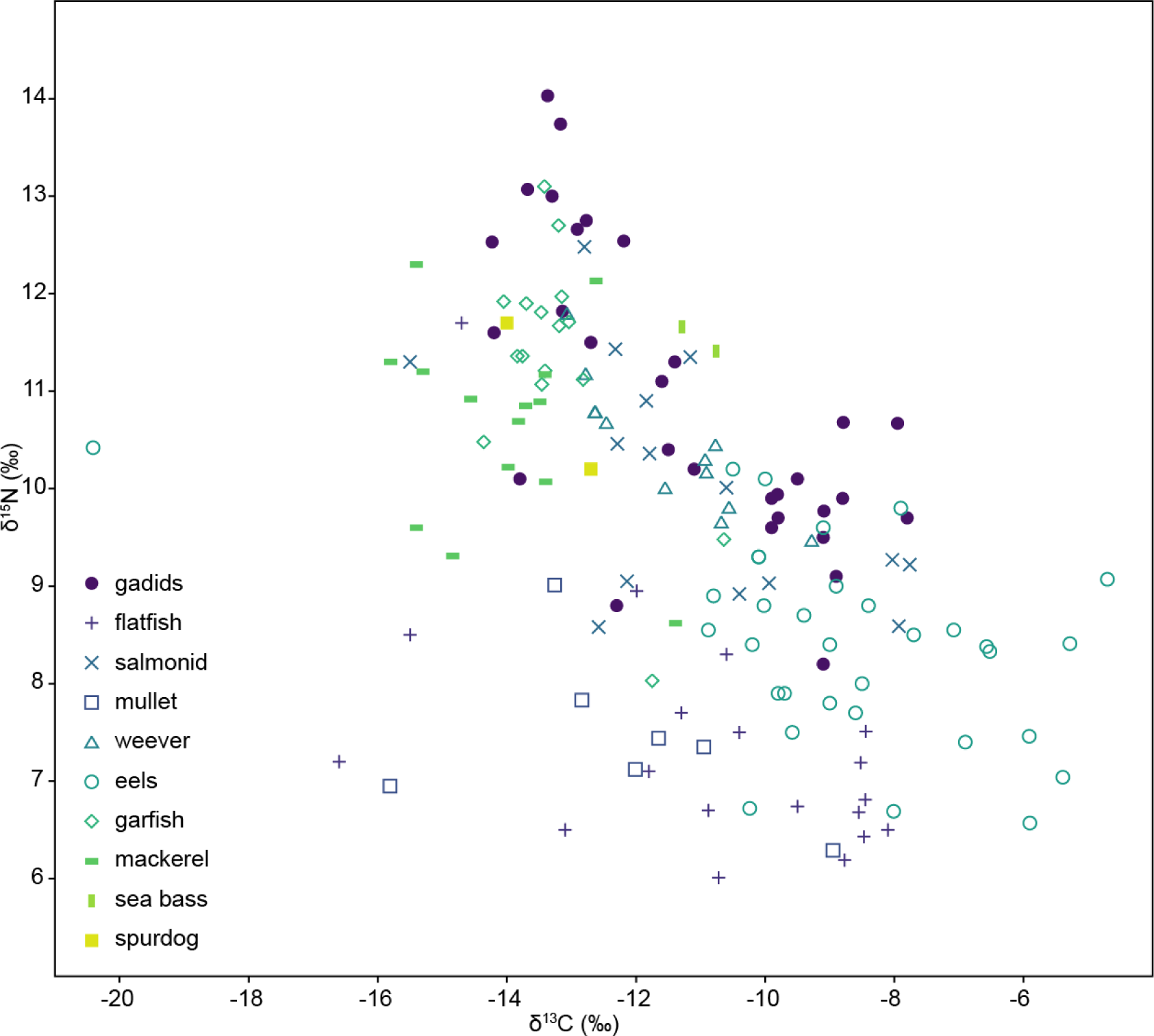
Collagen δ^13^C and δ^15^N values, marine fish taxa from prehistoric sites in coastal Denmark (Fischer et al. 2007a; Pleuger and Makarewicz 2020; Robson et al. 2012; Robson et al. 2016) (Supplementary Table 1).

Gadids (cod or haddock, but usually cod if determined to species) from eight sites show a significant negative correlation between δ^13^C and δ^15^N (Table 2). This pattern appears to reflect the role of eelgrass as a fish nursery: young cod consume invertebrates living in eelgrass (Hauser Jacobsen 2024; Lilley and Unsworth 2014), while mature cod live in deeper waters and consume increasing amounts of fish, including smaller cod, rather than invertebrates (Link and Garrison 2002). Thus cod δ^15^N values increase when they move from the eelgrass biotope to a higher trophic position within the pelagic food web, which is based on phytoplankton with lower δ^13^C values.

**Table 2:**
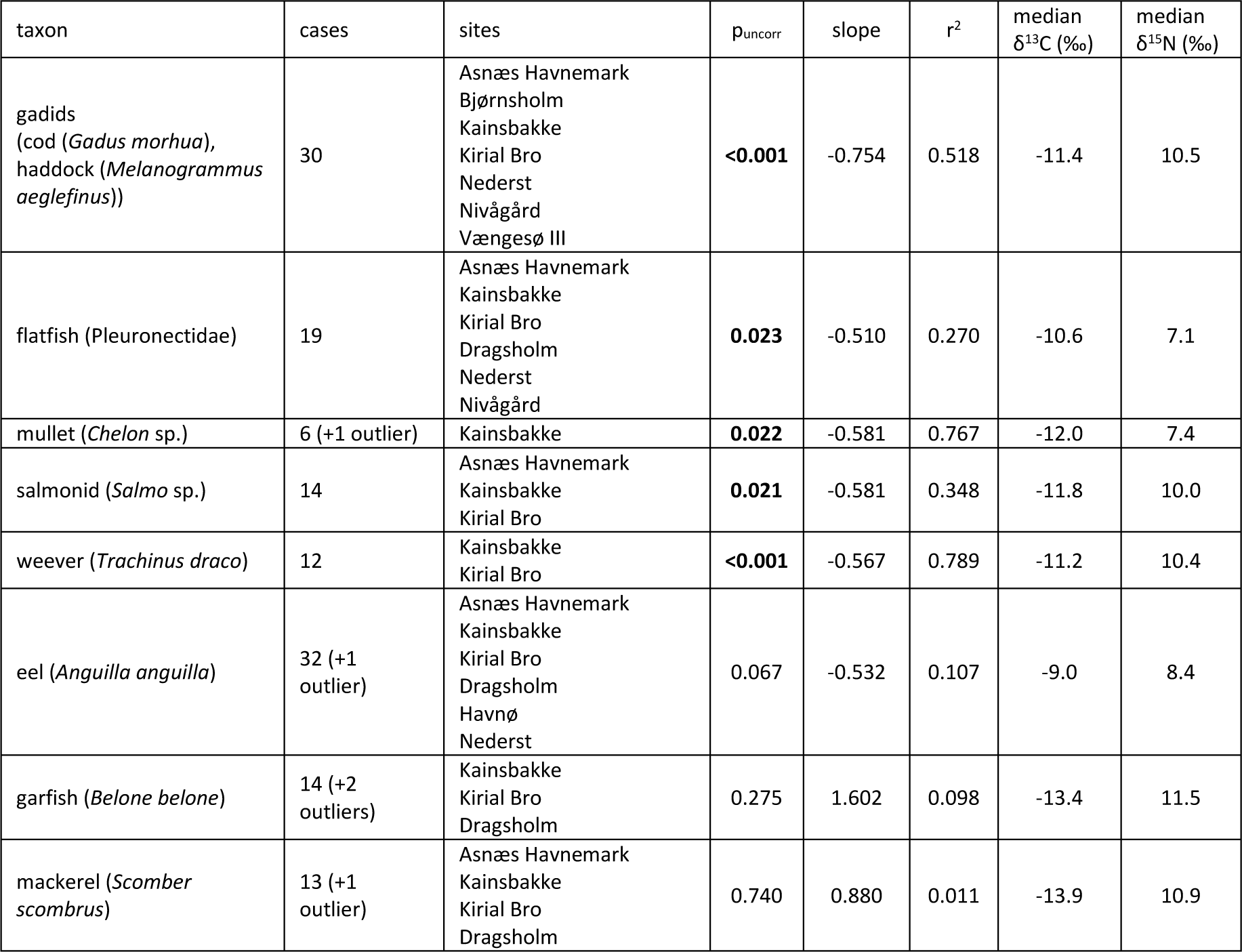
Reduced Major Axis regression between δ^13^C and δ^15^N among fish taxa from various prehistoric sites in Denmark (data in Supplementary Table 1). Significant correlations indicated in bold type. Outliers omitted from regressions may be anomalous due to habitat, differences in date, or perhaps misidentification.

| taxon | cases | sites | $p_{\text{uncorr}}$ | slope | $r^2$ | median $\delta^{13}\text{C}$ (‰) | median $\delta^{15}\text{N}$ (‰) |
| --- | --- | --- | --- | --- | --- | --- | --- |
| gadids<br>(cod (*Gadus morhua*),<br>haddock (*Melanogrammus*<br>*aeglefinus*)) | 30 | Asnæs Havnemark<br>Bjørnsholm<br>Kainsbakke<br>Kirial Bro<br>Nederst<br>Nivågård<br>Vængesø III | **<0.001** | -0.754 | 0.518 | -11.4 | 10.5 |
| flatfish (Pleuronectidae) | 19 | Asnæs Havnemark<br>Kainsbakke<br>Kirial Bro<br>Dragsholm<br>Nederst<br>Nivågård | **0.023** | -0.510 | 0.270 | -10.6 | 7.1 |
| mullet (*Chelon* sp.) | 6 (+1 outlier) | Kainsbakke | **0.022** | -0.581 | 0.767 | -12.0 | 7.4 |
| salmonid (*Salmo* sp.) | 14 | Asnæs Havnemark<br>Kainsbakke<br>Kirial Bro | **0.021** | -0.581 | 0.348 | -11.8 | 10.0 |
| weever (*Trachinus draco*) | 12 | Kainsbakke<br>Kirial Bro | **<0.001** | -0.567 | 0.789 | -11.2 | 10.4 |
| eel (*Anguilla anguilla*) | 32 (+1 outlier) | Asnæs Havnemark<br>Kainsbakke<br>Kirial Bro<br>Dragsholm<br>Havnø<br>Nederst | 0.067 | -0.532 | 0.107 | -9.0 | 8.4 |
| garfish (*Belone belone*) | 14 (+2 outliers) | Kainsbakke<br>Kirial Bro<br>Dragsholm | 0.275 | 1.602 | 0.098 | -13.4 | 11.5 |
| mackerel (*Scomber scombrus*) | 13 (+1 outlier) | Asnæs Havnemark<br>Kainsbakke<br>Kirial Bro<br>Dragsholm | 0.740 | 0.880 | 0.011 | -13.9 | 10.9 |

Size estimates are usually not reported in isotope studies. Enghoff (1994) charted estimated sizes of cod from 11 Ertebølle sites, and in all cases individuals >50cm in length are rare; this pattern has been confirmed subsequently (e.g. Price et al. 2018). In cod >50cm in length from medieval sites across northern Europe (smaller cod were not sampled), δ^13^C and δ^15^N are negatively correlated only in cod from the Kattegat/southwest Baltic (n=15, p_uncorr_ 0.043, slope -0.710, r^2^ 0.279, median δ^13^C - 13.1‰, median δ^15^N 12.5‰) (our analysis of data from Barrett et al. (2011)). Viking-era cod from Aarhus show a similar pattern (n=20, p_uncorr_ 0.001, slope -0.621, r^2^ 0.439, median δ^13^C -11.9‰, median δ^15^N 12.1‰) (our analysis of data from Swenson (2019)). The Danish medieval cod >50cm in length fall at the low-δ^13^C, high-δ^15^N end of the range in prehistoric cod, apparently confirming that the high-δ^13^C, low-δ^15^N end of this range represents smaller fish.

Several fish taxa in the Danish prehistoric material show similar isotopic patterns to cod, including flatfish, salmonids, weevers and mullets (Table 2), which suggests that within each taxon, isotopic variation is determined by the same mechanism. Individuals which fed mainly in inshore, eelgrass-dominated habitats have higher δ^13^C values; fish with higher δ^15^N values were presumably mature fish which fed mainly in offshore habitats where the food web was ultimately based on phytoplankton. This does not necessarily mean that mature fish were caught offshore, as they may have returned to the eelgrass zone, e.g. to spawn. Eels from six sites have the highest δ^13^C values (mostly -10 to -5‰), and must have fed mainly on invertebrates from the eelgrass biotope. Higher δ^13^C in eels appears to be associated with lower δ^15^N, but the data are slightly too scattered for this association to be significant.

All fish taxa with negative δ^13^C-δ^15^N correlations have relatively high median δ^13^C values (>-12‰, Table 2). Garfish and mackerel have much lower δ^13^C values (-16 to -13‰), indicating that they (and perhaps other low-δ^13^C taxa poorly represented in isotope data, such as spurdog and sea bass (*Dicentrarchus labrax*); Fig. 2) belonged to the pelagic food web, as at present (Muus and Nielsen 1998). Garfish and mackerel δ^13^C and δ^15^N values are not significantly correlated, but there is a positive trend in both species (Table 2). Herring (*Clupea harengus*) is not represented in the prehistoric isotope data, but in Viking-era material from Aarhus (Swenson 2019) it is clearly a pelagic species (median δ^13^C -14.6‰, n=12).

Seals with δ^13^C <-13‰ show a positive δ^13^C-δ^15^N correlation, also visible in early Mesolithic marine mammals from the Skagerrak coast of Sweden (Boethius and Ahlström 2018) (Fig. 3). This trend probably reflects trophic position within pelagic food webs, due to e.g. differential consumption of lower-trophic-level species such as sand eels (Ammodytidae spp.), and higher-trophic-level fish such as mature cod. Higher δ^13^C values in some Danish marine mammals (>-12‰) are associated with lower δ^15^N values, suggesting that some marine mammals also relied on eelgrass fish. Given their wide range of δ^13^C values, otters appear to have relied either on the eelgrass biotope, or lower-trophic-level fish in the pelagic food web (Fig. 3).

**Figure 3:**
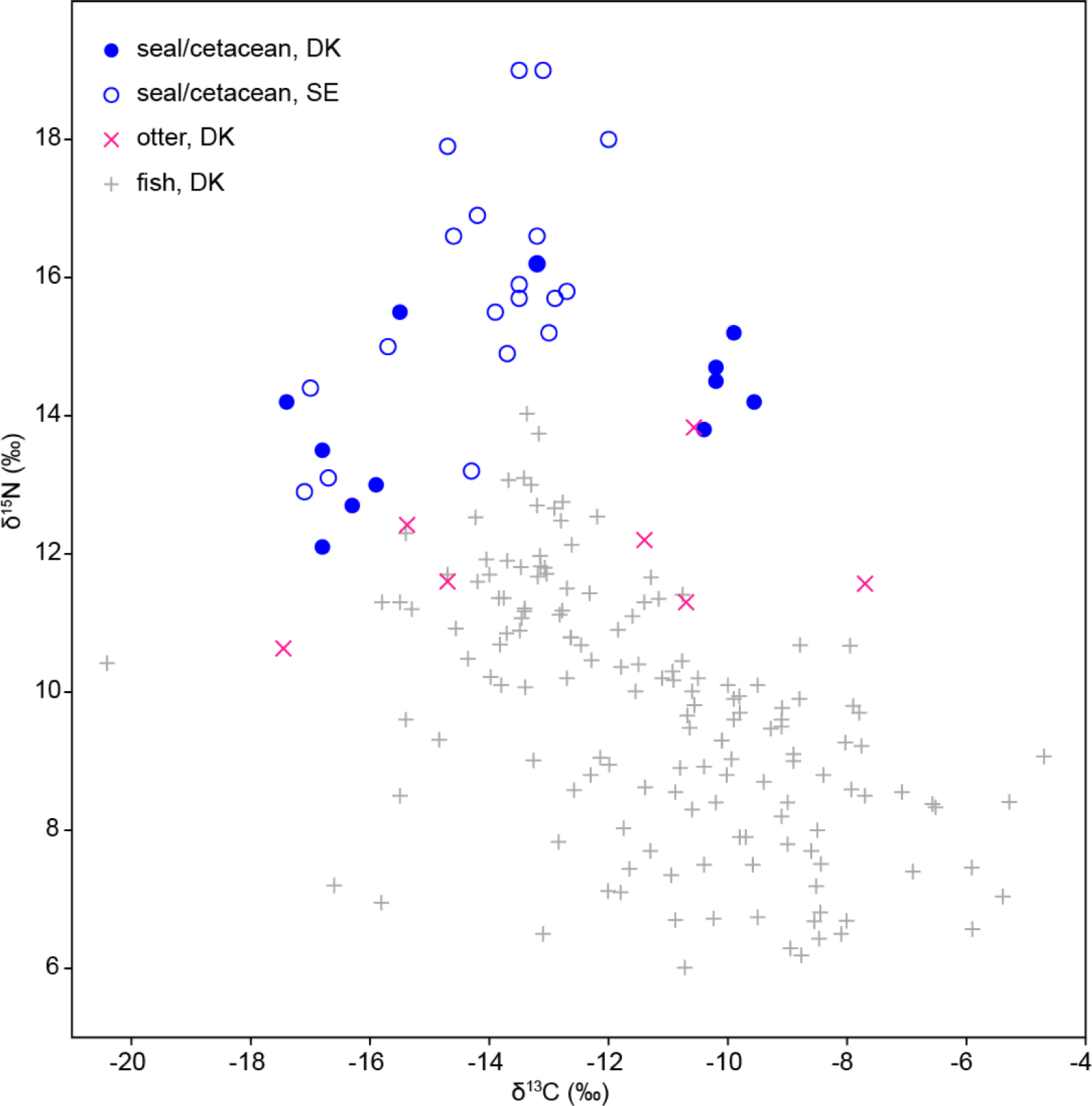
Marine mammal collagen δ^13^C and δ^15^N values from mid-Holocene Danish sites (Craig et al. 2006; Fischer et al. 2021; Fischer et al. 2007b; Robson et al. 2016) and early Holocene sites on the west coast of Sweden (Boethius and Ahlström 2018). Otter results from Table 1 and (Fischer et al. 2007b; Philippsen 2023). Danish prehistoric marine fish data (Fig. 2) plotted for comparison.

Marine-species δ^34^S data for prehistoric Denmark are limited, but available results (Fig. 4) show a negative correlation between δ^13^C and δ^34^S (p_uncorr_ 0.020, slope -1.075, r^2^ 0.561), consistent with the incorporation of eelgrass detritus into eelgrass-biotope food webs. Guiry et al. (2021b) relied on a negative correlation between δ^13^C and δ^34^S to argue that variation in the importance of seagrass habitats contributed to isotopic variation in 1800s fish bones from the Gulf of Mexico. In the past, fish or seal δ^34^S values <10‰ were attributed to the freshwater input to the Baltic (Craig et al. 2006; Fornander et al. 2008; Nehlich et al. 2013), implicitly assuming an unrealistically high concentration of low-δ^34^S sulfate in freshwater^1^. However, Craig et al. (2006) also observed a negative correlation between δ^13^C and δ^34^S values in seals and dogs from Bjørnsholm and Norsminde, and surmised that δ^34^S values were influenced by sulfate-reducing bacteria, without proposing a mechanism.

**Figure 4.**
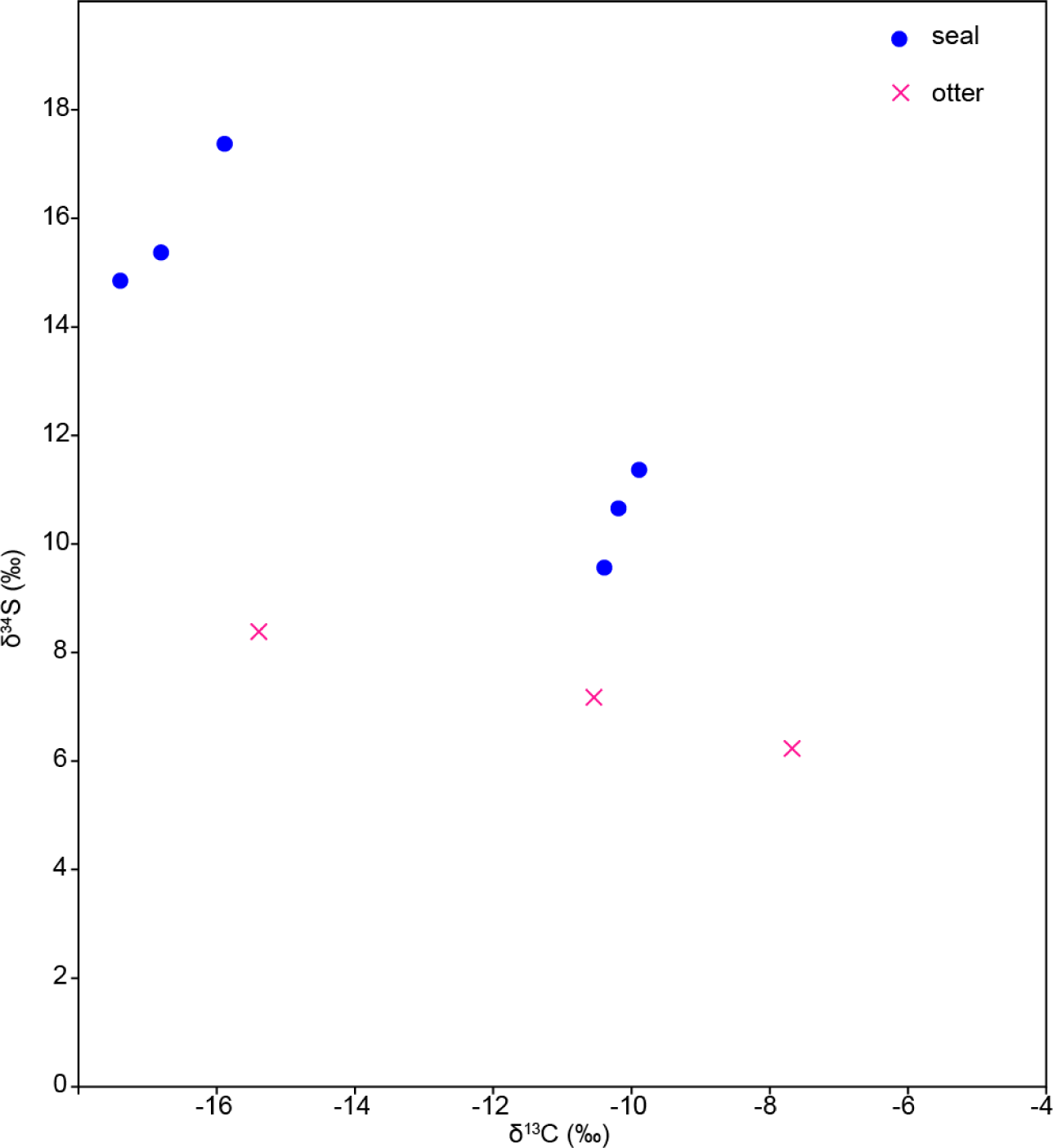
δ^13^C and δ^34^S in marine species from Danish prehistoric sites (this paper; Craig et al. (2006)).

#### 3.2.2. Dogs

Including our current results, δ^13^C and δ^15^N values are now available for 80 Danish ‘Ertebølle dogs’, almost half of which have been dated directly (Supplementary Table 2). Three published samples gave what now appear to be anomalous values, however, given the overall pattern of results (Fig. 5). A mandible from Syltholm (δ^13^C -22.4‰, δ^15^N 5.3‰ (Gron et al. 2024)) might belong to a herbivore – or perhaps a wolf (Ewersen et al. 2018) – while two specimens could be from seals: an unspecified bone from Ringkloster (not visible in Fig. 5; δ^13^C -16.6‰, δ^15^N 16.2‰ (Fischer et al. 2007a); species attribution from Rasmussen (1998)) and perhaps a canine tooth from Norsminde (δ^13^C -8.7‰, δ^15^N 14.1‰ (Craig et al. 2006)). As we cannot confirm their species determinations, we omit these three samples from our discussion, in case they were misidentified.

**Figure 5:**
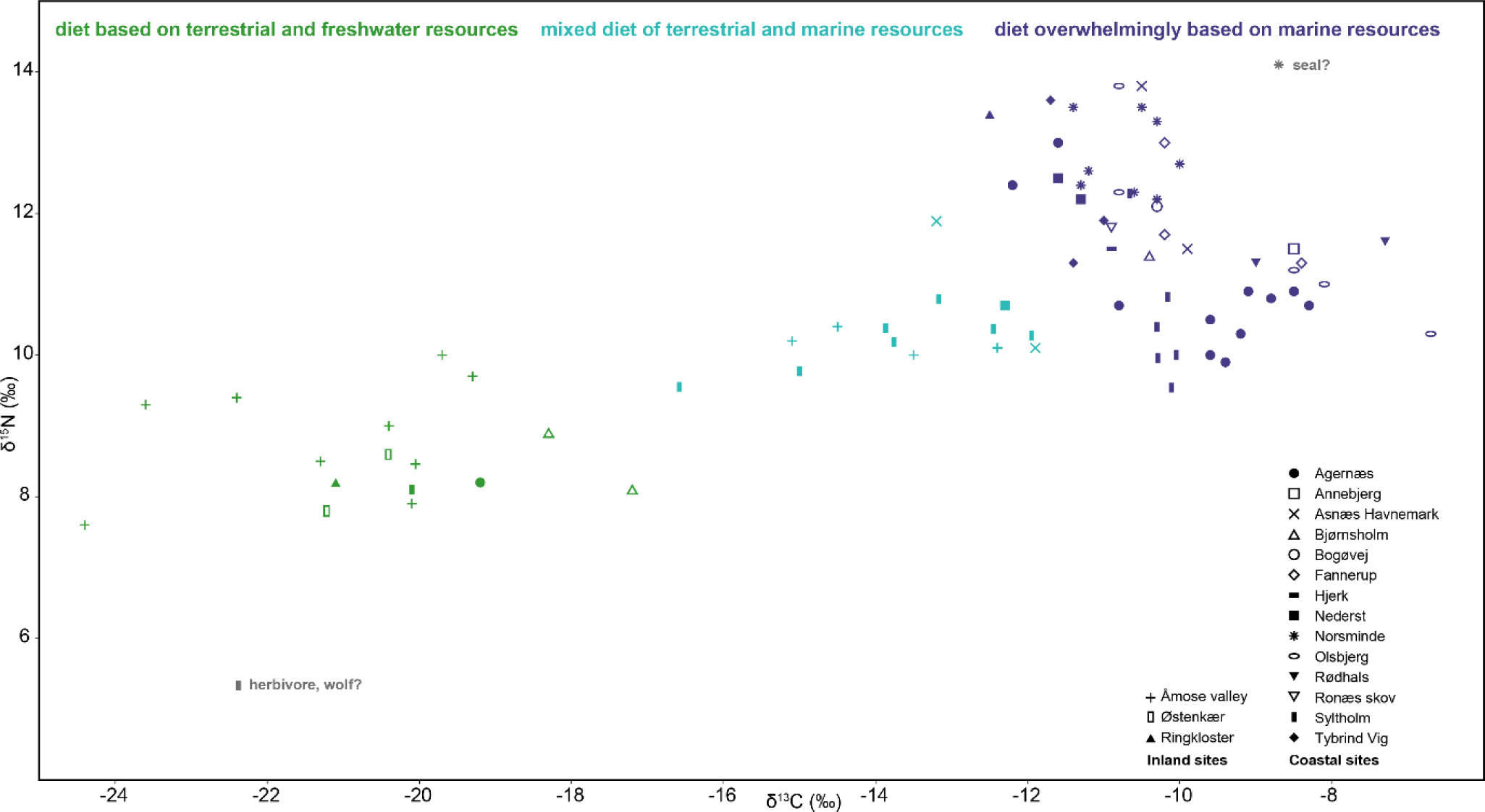
Dog collagen δ^13^C and δ^15^N values from Danish EBC sites, indicated by symbols (see legend). Data from Table 1 and (Craig et al. 2006; Fischer et al. 2021; Fischer et al. 2007a; Gron et al. 2024; Maring et al. 2024; Maring and Riede 2019; Philippsen 2023; Richter and Noe-Nygaard 2003). Colours reflect k-means clustering (k=3) of isotope values, omitting individuals with anomalous isotope values that we suspect may have been misidentified (grey).

Omitting these three cases, k-means clustering separates:

- 16 dogs, mostly found at inland sites in the Åmose valley, with isotope values (median δ^13^C - 20.3‰, median δ^15^N 8.5‰) indicating diets based mainly on terrestrial species and freshwater fish
- 14 dogs with intermediate isotope values (median δ^13^C -13.4‰, median δ^15^N 10.2‰), with mixed terrestrial-marine diets; these were found mainly in the inland Åmose valley, where they suggest seasonal mobility of Ertebølle groups (Fischer et al. 2007a), or at coastal Syltholm (Gron et al. 2024; Philippsen 2023), where some may date to the early Neolithic
- 47 dogs, found mainly at EBC coastal sites, with diets based primarily or overwhelmingly on marine resources (median δ^13^C -10.3‰, median δ^15^N 11.6‰).

The overwhelmingly marine diets of the third group are confirmed by a negative correlation between δ^13^C and δ^15^N values (p_uncorr_ <<0.001, slope =-0.940, r^2^ =0.300), as in many fully marine species. This correlation, reflecting differential reliance on the two marine biotopes, would be obscured if terrestrial and/or freshwater resources were important components of dog diets in this group.

Dog δ^34^S values confirm that δ^13^C and δ^15^N variation reflects the relative importance of pelagic and eelgrass biotopes. All δ^34^S values are well below those of oceanic sulfate, and in 13 dogs with overwhelmingly marine diets, from three sites^2^, δ^34^S is correlated with both δ^13^C (p_uncorr_ =0.006, slope -1.974, r^2^ = 0.515) and δ^15^N (p_uncorr_ =0.004, slope 0.417, r^2^ =0.537). Higher δ^34^S is associated with lower δ^13^C and higher δ^15^N (i.e. more pelagic-derived diets), while lower δ^34^S values are associated with higher δ^13^C and lower δ^15^N (more eelgrass-derived diets) (Fig. 6).

**Figure 6:**
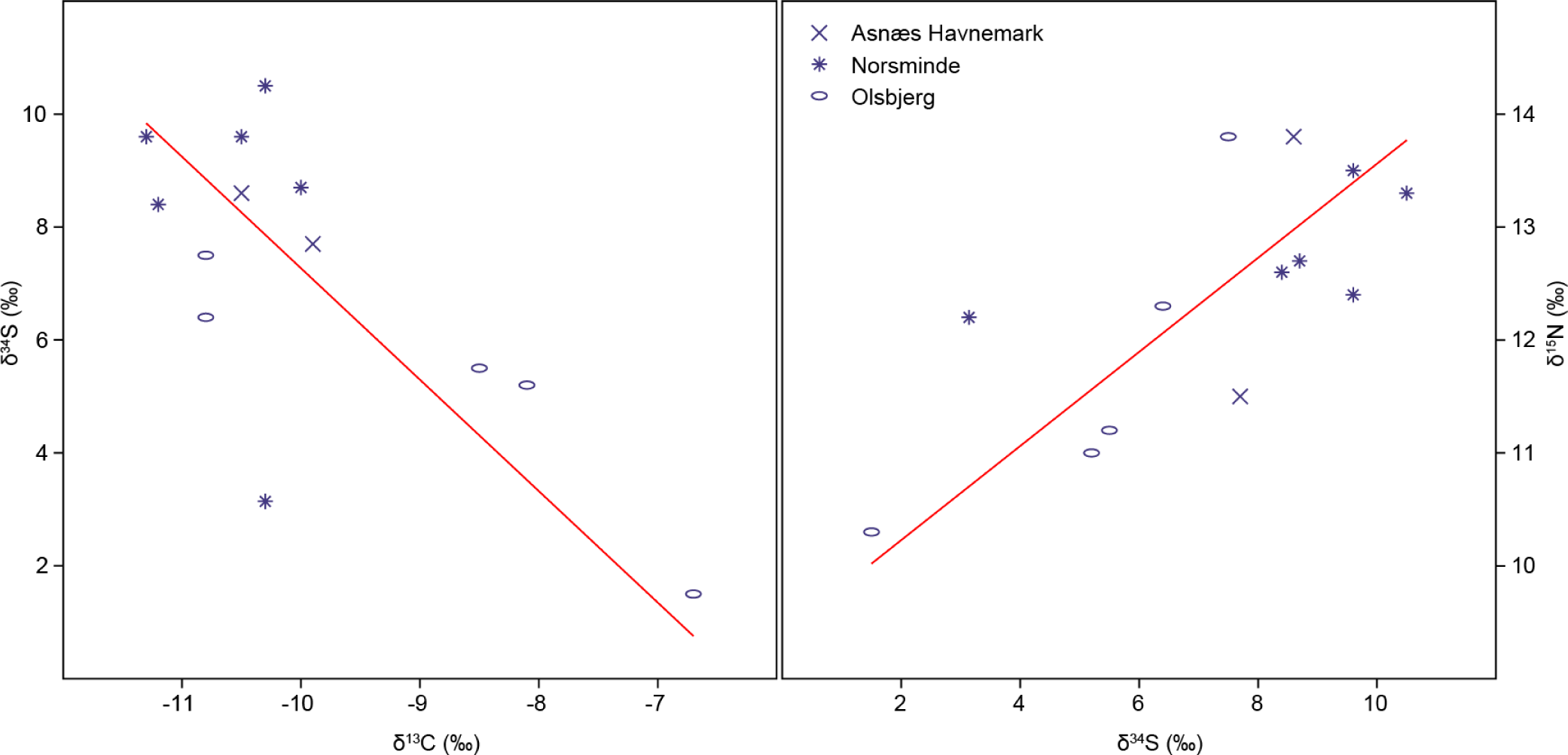
Dog collagen δ^34^S, δ^13^C and δ^15^N values from Danish EBC sites, and RMA regression lines. Data from Table 1 and Craig et al. (2006).

#### 3.2.3. Humans

It has long been known that human collagen δ^13^C values increased over the course of the Danish Mesolithic. Price and Gebauer (2005) suggested that human diets were only dominated by marine resources after ∼7450 cal BP, whereas Petersen (2015) argued that higher δ^13^C values in the Ertebølle epoch were due to increasing salinity, rather than increasingly marine diets. A long-term trend in Mesolithic human δ^13^C is clearly visible (Fig. 7), although in the earlier Mesolithic (Maglemose and Kongemose epochs), this may simply reflect gradually rising sea level, with the oldest sites on average further inland and having less access to marine resources of any kind. Earlier Mesolithic δ^13^C values (n=17) are positively correlated with δ^15^N (p_uncorr_ =0.004, slope 0.714, r^2^ =0.439), fitting the traditional model of two isotopically distinct food sources, terrestrial and marine biotopes. We see a sharp shift in δ^13^C values at ∼7400 cal BP, corresponding to the start of the EBC: previously, δ^13^C values > -14‰ are unknown; afterwards, almost 70% of individuals^3^ have δ^13^C between -13‰ and - 10‰ (Supplementary Table 3).

**Figure 7:**
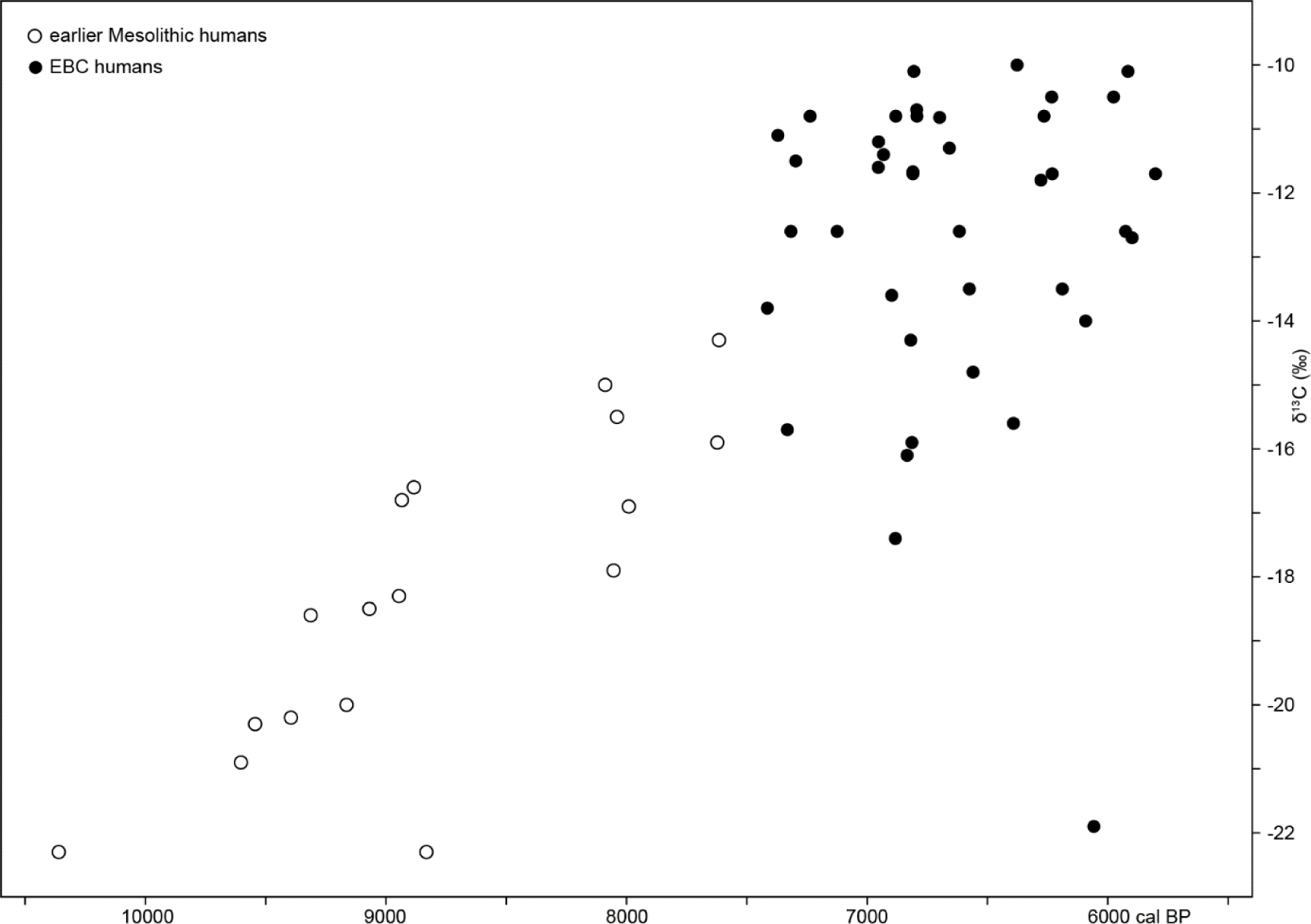
Danish Mesolithic human collagen δ^13^C versus median calibrated ^14^C age, after DRE correction (3.3.3). Data from (Allentoft et al. 2024; Allentoft et al. 2022; Fischer et al. 2007a; Price et al. 2007).

As with the dogs, k-means clustering of δ^13^C and δ^15^N values (Fig. 8) separates three groups:

**Figure 8:**
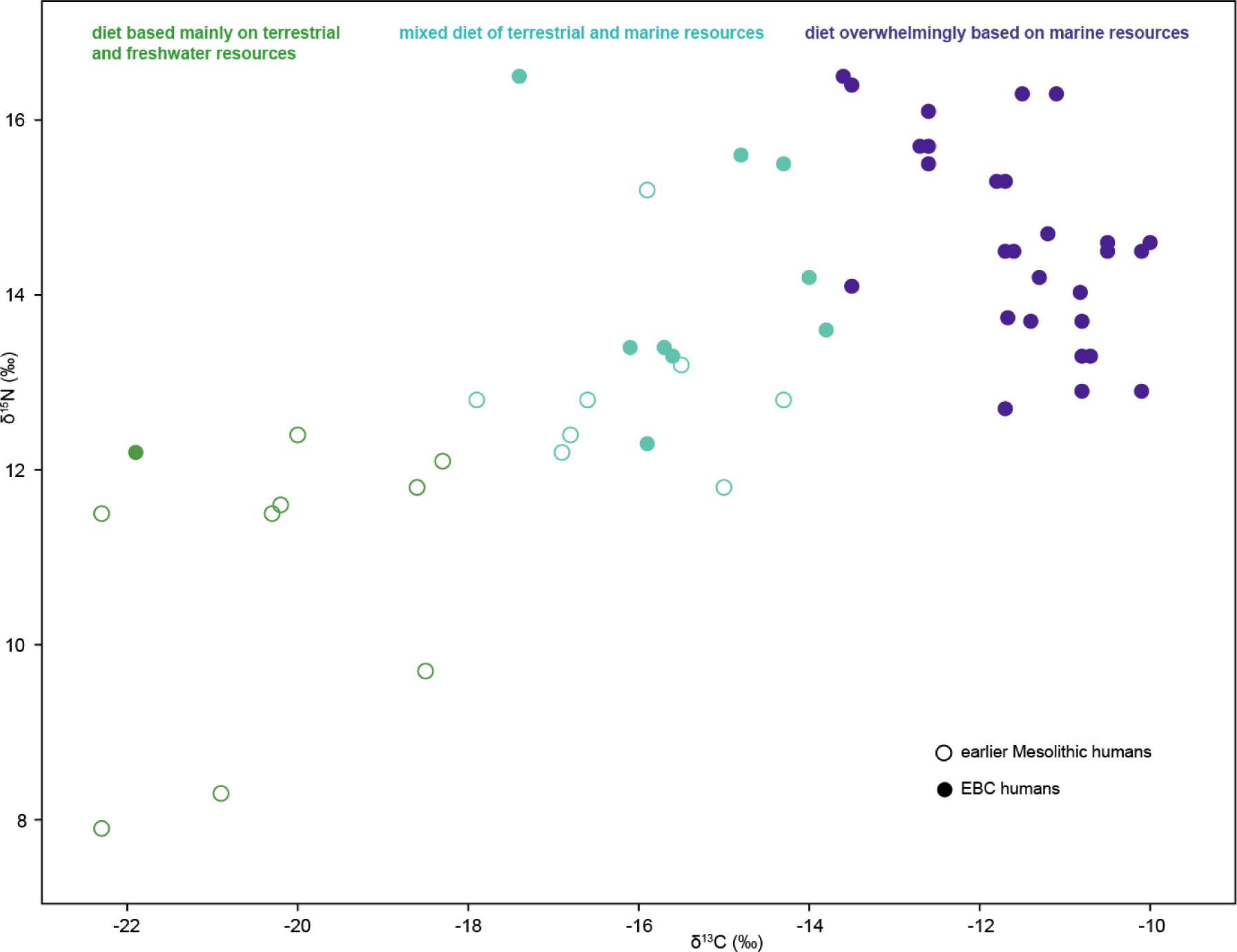
Danish Mesolithic human collagen δ^13^C and δ^15^N. Data from (Allentoft et al. 2024; Allentoft et al. 2022; Fischer et al. 2007a; Price et al. 2007). Colours reflect k-means clustering (k=3) of isotope values.

- 10 individuals with isotope values indicating diets based mainly on terrestrial species and freshwater fish, of whom 9 date to the earlier Mesolithic and only 1 to the EBC (median δ^13^C - 20.3‰, median δ^15^N 11.6‰)
- 17 individuals with intermediate δ^13^C values, indicative of mixed diets based on both terrestrial and marine resources; 8 of these individuals date to the earlier Mesolithic and 9 to the EBC (median δ^13^C -15.6‰, median δ^15^N 13.4‰)
- 28 individuals with diets mainly or overwhelmingly marine diets, all dated to the EBC (median δ^13^C -11.6‰, median δ^15^N 14.5‰).

As with the dogs, a negative δ^13^C-δ^15^N correlation in the third group (p_uncorr_ =0.002, slope =-1.103, r^2^ =0.314) suggests that differences in reliance on pelagic and eelgrass biotopes, rather than in the use of terrestrial resources, are the main cause of isotopic variation in this group, which represents 74% of dated EBC individuals. Human δ^15^N values are typically 1-2‰ higher than dog δ^15^N values for the same δ^13^C, either due to metabolic differences, or to trophic-level differences (e.g. if dogs ate more molluscs than humans, or humans sometimes ate dogs).

Only one EBC human bone δ^34^S value has been published, so we cannot look for correlations between δ^34^S and other isotopes. The authors (Gron et al. 2023) noted that this δ^34^S value was far lower than expected for a diet based on marine resources, and (following (Guiry et al. 2021a; Guiry et al. 2021b)) suggested ‘consumption of sulfide-derived resources’. We concur; in this context, sulfide-derived means ultimately derived from eelgrass biomass.

#### 3.2.4. Summary of isotopic patterns

We can account for observed patterns in δ^13^C, δ^15^N and even δ^34^S values in Danish marine vertebrates by recognising that two isotopically distinct primary producers, phytoplankton and eelgrass, explain much of the isotopic variation between individuals. This isotopic continuum in marine vertebrates is reflected in δ^13^C and δ^15^N data from the vast majority of EBC consumers (dogs and humans).

Salinity indirectly affects δ^13^C in marine animals. Archaeological fish bones from the southwest Baltic and Kattegat have higher δ^13^C values than fish bones from the Baltic proper, a phenomenon often attributed to higher salinity (Eriksson and Lidén 2002; Orton et al. 2011; Petersen 2015; Robson et al. 2016), because freshwater entering the Baltic has more negative DIC δ^13^C than oceanic water. Following Filipsson et al. (2017), DIC δ^13^C should be ∼2‰ higher in the Kattegat (22‰ salinity) than in the Baltic proper (8-10‰ salinity). This would affect δ^13^C in phytoplankton (and thus in the pelagic food web), which explains why modern herring δ^13^C values are ∼2‰ higher in the Kattegat than in the Baltic proper (Angerbjörn et al. 2006). Mature medieval cod δ^13^C values (Barrett et al. 2011) are 3-4‰ higher in the Kattegat/southwest Baltic than in the east Baltic, however, and juvenile modern cod δ^13^C values are ∼5‰ higher in the Kattegat than in the Baltic proper (Angerbjörn et al. 2006), reflecting the productivity of eelgrass in Danish waters and its importance as a fish nursery.

Much of the δ^13^C and δ^15^N variation at prehistoric Danish coastal sites can be attributed to variation in the contributions of pelagic and eelgrass biotopes. This is illustrated in Figure 9, which shows that:

**Figure 9.**
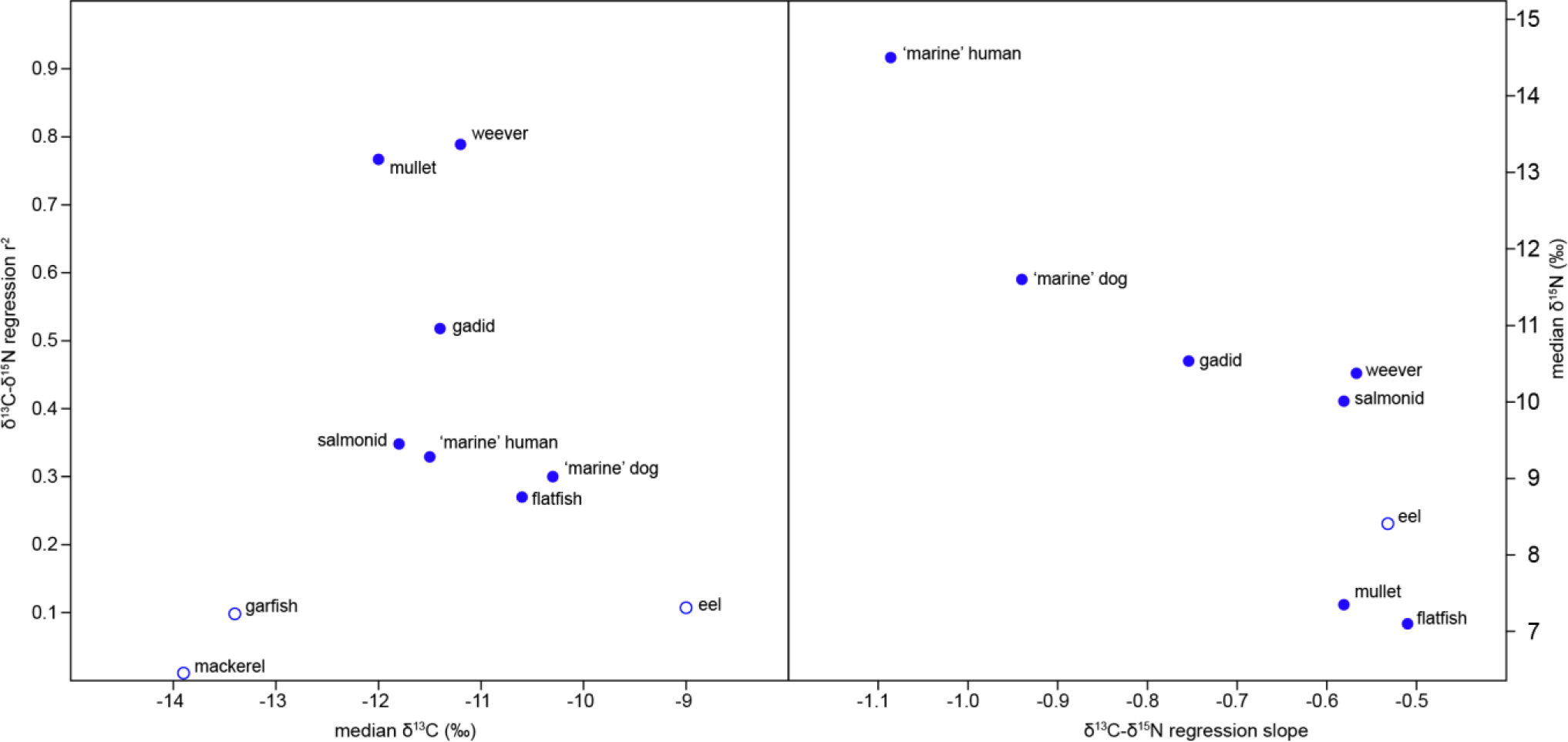
Median δ^13^C versus explanatory power (r^2^) and median δ^15^N versus effect (slope) of RMA regression of individual δ^13^C and δ^15^N values (Table 2, 3.2.2, 3.2.3). Filled symbols indicate that δ^13^C-δ^15^N correlations are statistically significant (p_uncorr_ <0.05).

- Significant negative δ^13^C-δ^15^N correlations occur only in taxa with intermediate δ^13^C values, i.e. in taxa depending to a significant degree on both the high-δ^13^C eelgrass biotope and the low-δ^13^C pelagic biotope. At either end of this continuum (mackerel, garfish, eel), <10% of δ^15^N variation is explained by δ^13^C variation, but 30-50% of the δ^15^N variation in cod, flatfish, salmonids, and dogs and humans with marine-focussed diets corresponds to δ^13^C variation
- The effect of these correlations is correlated with δ^15^N: the regression slope is steeper (more negative) for higher trophic level taxa, because the pelagic food chain was longer than the eelgrass food chain. Humans and dogs could have consumed marine mammals as well as larger pelagic fish, seals would have consumed mature cod as well as e.g. mackerel, herring and sand-eels, mature cod would have consumed smaller pelagic fish like herring but not seals, and so on.

Regression statistics for the Viking-age and medieval cod in the same region (3.2.1) fit the same patterns. Thus regression output is consistent and compatible with known ecological relationships between marine species, which suggests that available δ^13^C and δ^15^N data adequately describe marine food webs, and are not excessively distorted by stochastic factors or local baseline variation. The fact that the ‘marine’ dog and human clusters fit the same model of δ^13^C and δ^15^N variation as cod and flatfish confirms that these clusters represent essentially marine diets, in which consumption of terrestrial or freshwater resources had an almost negligible impact on isotope values.

### 3.3. Incorporating the eelgrass biotope in archaeological research

The contrasting isotopic signatures of eelgrass and phytoplankton should influence interpretation of isotope data from Danish coastal consumers (humans and dogs), including their ^14^C ages. Any archaeological application requires realistic assumptions, regarding:

- whether modern eelgrass productivity rates are applicable to the Ertebølle epoch
- parameterisation of food resources from eelgrass and pelagic biotopes (isotope values, energy and protein contents, associated uncertainties)
- ^14^C reservoir effects associated with eelgrass and pelagic biotopes.

#### 3.3.1. Assessing past eelgrass productivity

EBC human and faunal stable isotope data suggest that the eelgrass biotope was more productive and/or more readily exploited than other biotopes, but this may partly reflect the prominence of coastal kitchen middens in archaeological research, or the suitability for isotopic analysis of bones from kitchen middens. Past eelgrass productivity has never been investigated, but is relevant to the potential scale and resilience of the EBC economy, particularly if recovery biases might exaggerate the role of eelgrass in EBC subsistence.

Various proxy data have been used to reconstruct the marine environment during the Ertebølle epoch. Lewis et al. (2016) used diatom and mollusc evidence from four kitchen middens on the Kattegat coast to argue that salinity was close to modern conditions throughout the Ertebølle epoch, rejecting Rowley-Conwy (1984)’s idea that the scarcity of oysters in later levels of some EBC middens might reflect declining salinity. In the Little Belt, salinity was higher during the later 7th and earlier 6th millennia cal BP than it is now (Kotthoff et al. 2017). Lewis et al. (2020) also argued that seawater in the Danish archipelago was more saline in the Ertebølle epoch and subsequently than it is today.

Lewis et al. (2020) reported time-series from three sites of sedimentary pigment flux, a proxy for marine primary productivity in general (both phytoplankton and eelgrass). These time-series (which apparently track local salinity proxies) support the argument that the marine environment overall was highly productive during and beyond the Ertebølle epoch. The relative productivity of eelgrass and phytoplankton should be reflected in bulk organic δ^13^C from marine sediment archives, but to our knowledge, the only mid-Holocene organic sediment δ^13^C time-series is from the Limfjord, where it was used to compare inputs from marine and terrestrial sources (Philippsen et al. 2013).

Eelgrass itself is not particularly sensitive to temperature fluctuations (1.2.1), but other organisms in the eelgrass biotope may have been. Sea-surface temperature proxies in a core from the northeast Skagerrak (Rohde Krossa et al. 2017) appear to show exceptionally warm water during the Ertebølle epoch, and a sharp temperature drop towards its end (interpreted as increased outflow from the Baltic, which should also be visible in salinity proxies). By contrast, a core from the Gotland basin suggests that the Baltic sea-surface temperature was relatively cold throughout the Ertebølle epoch, but then rose sharply (Warden et al. 2017). We are not convinced by these contradictory patterns in water temperature proxies, and suggest that eelgrass growing conditions in Danish waters were at least as favourable during the Ertebølle epoch as in the recent past.

#### 3.3.2. Human diet reconstruction

Ertebølle human and dog δ^13^C and δ^15^N values have been used to estimate the contributions to diet of marine, terrestrial and freshwater resources, and thus to infer mobility between coastal and inland sites (Fischer et al. 2007a; Maring and Riede 2019). The vast majority of Ertebølle human and dog diets appear to have been dominated by marine resources (3.2.2, 3.2.3), however. To quantify diet variation among EBC individuals, it is more important to treat pelagic and eelgrass biotopes as isotopically distinct sources of marine foods.

To illustrate the importance of the eelgrass biotope to human diets in the Ertebølle period, we use an unrouted (concentration-independent) FRUITS model (Fernandes et al. 2014) to estimate the contributions of terrestrial, pelagic and eelgrass biotopes to the diets of the 38 individuals dating after 7500 cal BP in Fig. 7 (details in Appendix B). An unrouted model estimates each source’s contribution to dietary protein intake only, but this is equivalent to its share of total diet if the energy density of each source is similar (Fernandes 2016). As both pelagic and eelgrass biotopes include energy-dense species (e.g. seals and eels), and terrestrial resources include wild plant foods (e.g. hazelnuts), this may be realistic. The FRUITS estimates suggest that in most cases, >80% of dietary protein was from marine sources, and that the eelgrass biotope was usually more important than the pelagic biotope (Supplementary Table 4); on average, the eelgrass biotope accounted for 46% of overall human diet. No chronological trends are visible in the FRUITS output: there is no indication that Ertebølle FHG diets were increasingly based on terrestrial, pelagic or eelgrass resources, or that they became more or less specialised over time (Fig. 10).

**Figure 10:**
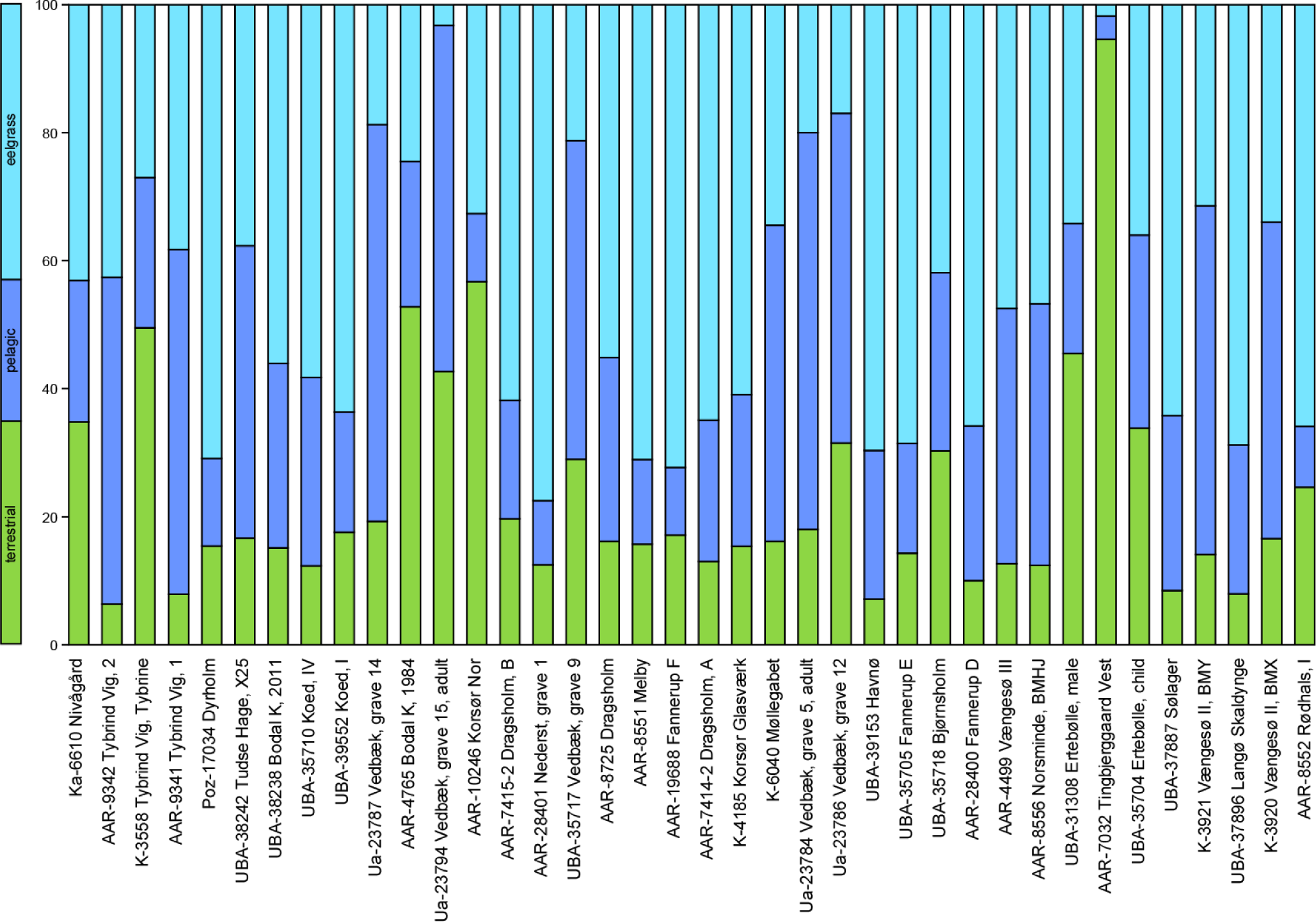
Stacked median estimates of the percentage contribution of terrestrial, pelagic and eelgrass biotope resources to EBC individual diets, based on human δ^13^C and δ^15^N values. Cases are ordered from left to right according to median date, after correction for marine DREs (3.3.3). Our model probably under-estimates the DRE in the Tingbjerggård individual, whose diet would have contained freshwater fish; this individual therefore would date somewhat later.

#### 3.3.3. DRE correction in human bones

##### Marine Reservoir Effects (MREs)

The slow circulation and upwelling of deep ocean water leads to an apparent age of several hundred years in surface water DIC. This marine reservoir effect (MRE) is transferred through food webs, affecting ^14^C ages of consumers, including humans and dogs. Global average MRE may be adjusted by a ΔR value, reflecting local ocean circulation patterns. The ^14^C ages of nine known-date, pre-1950 marine molluscs from Danish coastal sites facing the Kattegat (Heier-Nielsen et al. 1995) suggest a ΔR relative to the Marine20 calibration model (Heaton et al. 2020) of -140±84y (http://calib.org/marine/, accessed 4 Nov 2023). If this ΔR was valid in the past, the local MRE at 6000 cal BP, when global MRE was ∼550y (Heaton et al. 2020), would have been ∼400y.

Early Neolithic oyster shells from Nekselø had an MRE of only 273±18y, however (Fischer and Olsen 2021). Oysters presumably consumed a mixture of detritus derived from eelgrass, epiphytes and pelagic phytoplankton, but their shells would have been formed directly from DIC, and should provide a realistic ^14^C signal for DIC in shallow coastal water. The contribution of freshwater DIC at estuarine sites is a potential complication (Olsen et al. 2017; Olsen et al. 2009), but the existence of oyster beds implies relatively saline conditions, which also favoured eelgrass productivity, so we regard oyster shell MREs as good proxies for eelgrass MREs. Using Bayesian chronological models (Appendix C), we calculate ΔR relative to Marine20 in marine shells at four Ertebølle sites, Asnæs Havnemark (Price et al. 2018), Hjarnø (Astrup et al. 2021), Horsens Fjord and Tempelkrog (Olsen et al. 2017; Olsen et al. 2009). Our ΔR estimates range from -380±50y (Horsens Fjord 988-818cm) to - 170±55y (Asnæs Havnemark), suggesting that Nekselø (ΔR -234±61y) could be fairly typical. For simplicity, in calculating DRE corrections (3.3.3) we apply the Nekselø ΔR of -234±61y to eelgrass food-sources, just as our FRUITS model (3.3.2) applies the same food-source δ^13^C and δ^15^N values to each biotope throughout Denmark.

Considering the outflow of brackish water from the Baltic, the global surface ocean calibration model Marine20 (Heaton et al. 2020) may exaggerate MREs in pelagic species in the Kattegat, but a more accurate model requires an estimate both of the extent to which oceanic water was diluted by Baltic outflow during the Ertebølle epoch, and of the DIC concentrations in oceanic and Baltic water. Empirical ^14^C data from tightly dated pelagic species are scarce. Whales caught in Norwegian waters in the late 1800s (Mangerud et al. 2006) had a ΔR of ∼-160y relative to Marine20, and similarly negative ΔR values apply to three historical cetaceans from the west coast of Sweden (Olsson 1980). The low ΔR of post-medieval cetaceans could be misleading, however, as stratified archaeological finds in Ireland and Scotland suggest that North Atlantic ΔR was ∼200y higher (more positive) in the mid-Holocene than in the medieval period (Ascough et al. 2009; Reimer et al. 2002). To calibrate the pelagic component of EBC diets, therefore, we simply apply Marine20, with a ΔR of 0±50y. This is not intended to be fully realistic, but allows us to visualise the potential impact on DRE corrections of differentiating pelagic and eelgrass biotopes.

##### Dietary Reservoir Effects (DREs)

It was traditionally assumed that a ‘fully marine’ diet (in which 100% of dietary protein came from marine sources) produced a characteristic δ^13^C value in human bone, based on analogies with Norse and indigenous diets in Greenland and North America (Arneborg et al. 1999; Chisholm et al. 1982), Danish skeletons often had δ^13^C values above the -12.5‰ used as the 100% marine endpoint in Greenland, so researchers assumed that the highest human δ^13^C represented a 100% marine diet, or used faunal isotope data to infer a δ^13^C of -10‰ in humans eating only marine species, and a δ^13^C of e.g. -21‰ corresponding to a 100% terrestrial diet (Fischer et al. 2007a). Individual DREs were then calculated by estimating the marine protein intake by interpolation between these two endpoints, and multiplying it by the applicable MRE (e.g. 400y or 273y (Fischer et al. 2024; Gron et al. 2023)).

Our 3-source diet reconstructions (3.3.2) provide more realistic estimates of the contributions of terrestrial, pelagic and eelgrass-derived carbon to individual ^14^C ages. These estimates, coupled with separate MRE estimates for pelagic and eelgrass resources, should lead to more accurate DRE corrections. Under our 3-source model, most Ertebølle DREs are somewhat smaller than under the traditional interpolation approach, so the true dates of these individuals will be slightly earlier than previously assumed. Figure 11 shows that using a 400y MRE, the interpolation approach over-estimates DREs in the most eelgrass-based diets by up to ∼120y, and underestimates DREs in the most pelagic-based diets by up to ∼120y, compared to our model. With a lower MRE (e.g. 273y), the interpolation approach would give more accurate DRE estimates for eelgrass-based diets, but risks significantly under-estimating DREs in cases with a high intake of pelagic species, even if ΔR in the pelagic biotope was similar to that in the eelgrass biotope (as the interpolation approach over-estimates the terrestrial share of diet in these cases).

**Figure 11:**
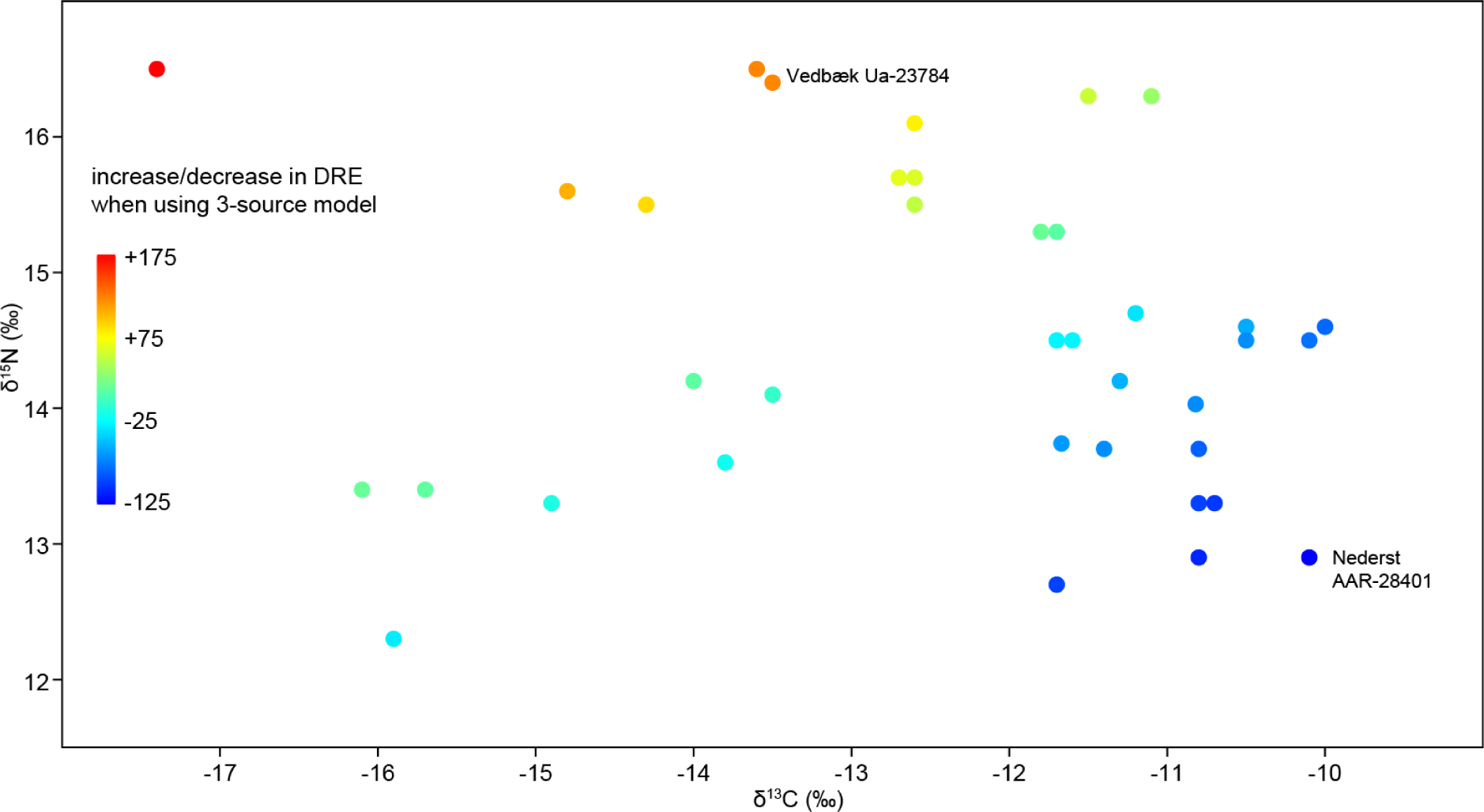
δ^1^3C and δ^15^N values of EBC humans, colour-coded by differences in DREs obtained using our 3-source model compared to the traditional δ^13^C interpolation approach with a 400y MRE (Supplementary Table 4). Two extreme cases are labelled.

To illustrate the potential consequences of our 3-source diet reconstruction, we consider two extreme cases (Fig. 11), dated by Allentoft et al. (2024). Regardless of the MRE employed, these two individuals are chronologically inseparable when DREs are estimated assuming a single marine food group, but if the pelagic MRE was significantly greater than the eelgrass MRE, the Nederst individual may be 200-400 years older than the Vedbæk (Henriksholm-Bøgebakken) burial (Table 3, Fig. 12).

**Figure 12:**
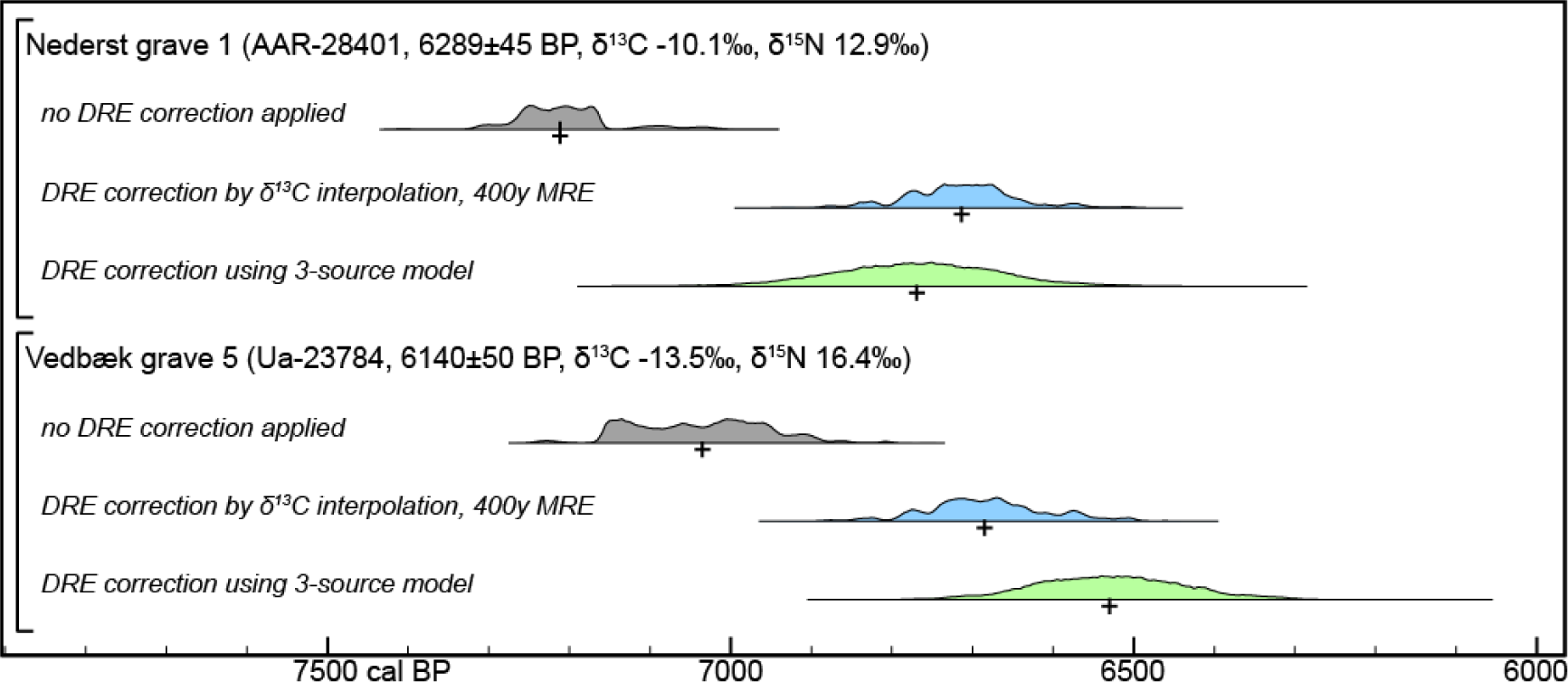
Calibrated dates of two EBC human bones using different DRE correction models. For Vedbæk grave 5, the median date (+) obtained with our 3-source model is ∼150y later than when using δ^13^C linear interpolation and a 400y MRE; for Nederst, the 3-source model dates the bone ∼60y earlier.

**Table 3.** Calibration of ^14^C ages for two humans of EBC association. Blue cell shading: previously applied 2- δ^13^C end-member interpolation model. Green shading: the 3-source model presented in this paper.

| case | lab code | $\delta^{13}\text{C}$ (‰) | $\delta^{15}\text{N}$ (‰) | $^{14}\text{C}$ age (BP) | median date (cal BP), no DRE correction | interpolation diet and DRE (100% marine $\delta^{13}\text{C}$ -10‰, MRE 400y) | median date (cal BP), interpolation DRE correction | 3-source model mean diet and implied DRE | median date (cal BP) 3-source model |
| --- | --- | --- | --- | --- | --- | --- | --- | --- | --- |
| Nederst grave 1 | AAR-28401 | -10.1 | 12.9 | 6289±45 | 7212 | 99% marine, DRE 396y | 6714 | 12% terrestrial, 11% pelagic, 76% eelgrass; DRE 270y | 6771 |
| Vedbæk grave 5 | Ua-23784 | -13.5 | 16.4 | 6140±50 | 7036 | 68% marine, DRE 273y | 6686 | 18% terrestrial, 62% pelagic, 20% eelgrass; DRE 395y | 6561 |

### 3.4. Areas for future research

#### 3.4.1. Model parameterisation

For illustrative purposes, we have estimated the diets and DRE-corrected ^14^C dates of 57 Mesolithic humans using the same model parameters (stable isotope and MRE signatures of pelagic and eelgrass biotope food sources) throughout the Danish archipelago and throughout the Mesolithic. Although our estimates are based on empirical data and are undoubtedly more realistic than those obtained by traditional interpolation between δ^13^C values for 100% terrestrial and 100% marine diets, for detailed case studies we recommend a more critical assessment of applicable parameter values. Eelgrass productivity will have varied geographically, partly as a function of salinity, affecting δ^13^C values in eelgrass biotope resources; this may explain, for example, why dog δ^13^C values at the eelgrass biotope end of the marine diet cluster (Fig. 5) are higher at Agernæs than at Syltholm, and even higher at Olsbjerg and Rødhals. Isotopically, the Syltholm marine dogs plot closest to the most marine dogs from Rosenhof, 50km away on the German coast (Fig. 1). The MRE applicable to mid-Holocene pelagic resources is difficult to measure empirically, but could be constrained if marine cores dated by ^14^C ages of foraminifera were tightly correlated with well-dated terrestrial proxy records (e.g. micro-tephra, pollen influx). It is tempting to recommend using δ^34^S as an additional dietary proxy, as low δ^34^S values are clearly associated with the eelgrass biotope, but massive fractionation by sulfate-reducing bacteria leads to a wide range (>20‰) in eelgrass δ^34^S values (Frederiksen et al. 2006). Terrestrial herbivores in coastal zones may also have had anomalously high δ^34^S values due to sea-spray effects. Local studies on well-preserved fish and herbivore bones are required to provide meaningful estimates of δ^34^S in dietary protein consumed by humans and dogs, with strict quality control to ensure that faunal δ^34^S values are not affected by diagenesis.

#### 3.4.2. Ertebølle technology

The importance of the eelgrass biotope may be reflected in EBC technology, but its impact has not been investigated. EBC FHGs employed a wide range of fishing technologies, which were already used earlier in the Mesolithic (Hausmann et al. 2022). Canoes, nets, fishhooks and harpoons were available for offshore fishing and hunting (Andersen 1995; Andersen 1997), but some marine mammals were probably taken on the coast (seal hunting and cetacean strandings (Møhl 1970)). Stationary capture systems (baskets, fences, weirs) were designed for use in shallow water, as were fish spears and leisters for taking eels, etc. The same technologies were (to some extent) also employed on rivers and lakes (Fischer 2007). Thus Ertebølle groups were well-equipped to exploit the eelgrass biotope, and their fishing technology was diverse and adaptable.

The Ertebølle epoch is notable for the adoption of pottery, which could have implications for the subsistence economy. Pointed-base cooking pots and shallow bowls, known as ‘lamps’, were widely distributed only in the later part of this epoch. It is not clear why pottery was adopted in ∼6500 cal BP, and not several centuries earlier, as typical diets do not appear to have changed over the course of the Ertebølle epoch (Fig. 10). Organic residue analysis provides insights into what was cooked in pointed-base cooking pots, both through lipid biomarkers and through bulk food-crust and lipid compound-specific isotope analyses; marine resources predominate (Lucquin et al. 2023). Much of the isotopic variation in organic residues must therefore reflect whether pelagic or eelgrass species were cooked, separately or together, but no attempt has been made to differentiate these sources. Lamps required a source of fat, which at coastal sites was most likely seal (mostly pelagic), eel (eelgrass biotope), or a mixture of both. Compound-specific δ^13^C values on lipids extracted from lamps from Danish sites (Robson et al. 2022) are often too high for the marine fat reference ellipse (which is based mainly on North Atlantic fish, seals and invertebrates), suggesting that the lamps were fuelled with fat from eelgrass resources, most likely from eels. Nevertheless, both lamps and pointed-base cooking pots were also widely used by FHGs in Eastern Europe, so we cannot attribute their adoption in the EBC to the productivity of eelgrass in Danish waters.

#### 3.4.3. Palaeodemography

The productivity of the eelgrass biotope, combined with EBC diet reconstructions, should provide insights into potential populations during the Ertebølle epoch. Based on Astrup (2018)’s reconstructions of past coastlines, by the mid-7th millennium cal BP there were ∼120,000 ha of seabed within 100 m of the Danish coast. Given modern NPP data (1.2.1), if half this area was suitable for eelgrass, it may have photosynthesised 0.4 million tonnes of carbon annually, theoretically enough to feed 2 million people, assuming an annual carbon budget of 200kg per capita, or 4 million, given that (on average) eelgrass resources provided only about half of human diets (3.3.2). According to Zhu et al. (2021), however, less than 0.01% of NPP is extracted by modern foragers in terrestrial biotopes, even in plant-based economies; in animal-based economies, forager offtake may be only 0.001% of NPP. A 0.001% offtake would imply a total EBC population in Denmark of only about 40 individuals.

Zhu et al. (2021) found that modern forager population densities in terrestrial biotopes increase according to a logarithmic function of NPP (following Hatton et al. (2015), we expect a similar rule to apply for aquatic biotopes). Applying this function to Danish eelgrass suggests a population density of 0.25 km^-2^, and a total population of 150 subsisting on 60,000 ha of eelgrass meadows, or twice that if terrestrial and pelagic resources are included. Modern forager population density data are dispersed, by up to a factor of ten at any NPP value (Zhu et al. 2021). Ertebølle population densities may have been relatively high, due to the spatial and seasonal predictability, and low inter-annual variability, of eelgrass resources. Compared to terrestrial subsistence strategies, however, trophic inefficiency (the loss of energy at each step in the trophic ladder) would reduce the available offtake (even eels and flatfish are at least one trophic level higher than forest herbivores), although exploiting the eelgrass biotope would have been more trophically efficient than depending on pelagic resources.

The extent and NPP of eelgrass meadows are also important variables. The extent of eelgrass in Danish waters was estimated as 6726 km^2^, or 672,600 ha, in 1914 (Boström et al. 2014), although much of this was in offshore waters; nevertheless, we could increase our Ertebølle population estimates by assuming that a larger area of eelgrass was exploited (e.g. that fish captured inshore also foraged in offshore eelgrass meadows). If humans effectively exploited an eelgrass area of 200,000 ha, whose NPP was relatively high (1000 gC m^2^ a^-1^), and if the offtake was 0.01% of NPP, we obtain a total EBC population in Denmark of ∼2000, if half their diets came from pelagic and terrestrial food webs. This is still surprisingly low, but represents a useful point of departure in discussing Ertebølle demography, and in considering how EBC capture and storage technology might have increased offtake from eelgrass biotopes above 0.01% of NPP. As Zhu et al. (2021)’s observed relationship between NPP and forager population density is logarithmic, small changes in NPP may have caused relatively large changes in population density (e.g. a 30% decline in NPP is predicted to result in a 60% decline in population density). This is particularly pertinent to the question of the end of the Ertebølle epoch.

#### 3.4.4. Resilience and fragility

Various studies have proposed environmentally deterministic models to account for the end of the Ertebølle epoch. When ^14^C dating and imported artefacts showed that the start of the Ertebølle epoch coincided with the start of farming in northern Germany, it became necessary to explain why Ertebølle fisher-hunter-gatherers did not adopt farming for over 1000 years, and why farming spread rapidly across the entire Danish archipelago when it eventually did (Rowley-Conwy 1984; Zvelebil and Rowley-Conwy 1984). For most of this period, it would have been possible to incorporate plant and animal domesticates into the subsistence strategy, but we see no evidence of this happening until perhaps the final century of the Ertebølle epoch (Lucquin et al. 2023). By then the Ertebølle population may have been in decline, given that there was little if any genetic contribution from Ertebølle FHGs to the subsequent Neolithic population of Denmark (Allentoft et al. 2024). A hypothetical (temporary) crash in eelgrass NPP in the early 6th millennium cal BP could explain the speed of the transition, but there is currently no convincing evidence of large sea-level fluctuations at this time, or of declines in salinity or water temperature in the Kattegat (3.3.1). Thus it is difficult to attribute the demise of the Ertebølle system to exogenous environmental change.

We have not investigated feedback mechanisms between Ertebølle subsistence strategies and the ecology of the eelgrass biotope. Over-exploitation might have damaged eelgrass-biotope fish stocks without affecting eelgrass primary productivity, and might be reflected in increasing use of species that were previously disregarded, or of smaller individuals of preferred species. There is no evidence of eels decreasing in size over time at the long-lived Havnø midden (Robson et al. 2013), but to reveal any overall trends, high-resolution archaeozoological evidence from multiple sites should be dated and collated. Another indication of resource stress could be greater dietary diversification. Among the dated human bones, we do not see that diets became more broad-based (or less dependent on eelgrass resources) towards the end of the EBC epoch (Fig. 10). Directly dated dog bones from the final centuries of the Ertebølle epoch are scarce, except at inland sites in the Åmose valley (Supplementary Table 2). Again, there is no obvious sign that the eelgrass-based FHG economy became environmentally unsustainable before the end of the Ertebølle epoch, but more detailed investigation might reveal evidence of fragility in its final centuries.

#### 3.4.5. Beyond EBC Denmark

There is a clear contrast between Early Neolithic human stable isotopes in Denmark, which imply overwhelmingly terrestrial diets, and fishing technology and pottery organic residue analyses, which show little change from the EBC focus on marine resources (Lucquin et al. 2023). A better understanding of marine resources might help to resolve this enigma. Early Neolithic diets were probably too terrestrial to reveal the contributions of pelagic and eelgrass biotopes to any marine component, but compound-specific δ^13^C of lipids in pottery, together with biomarker analysis and food-crust ^14^C, δ^13^C and δ^15^N values, should reflect the relative importance of eelgrass resources.

Because the primary productivity of eelgrass in the Kattegat and Belt Sea was much higher than in the Baltic proper, we expect different adaptations at EBC sites in southern Sweden and northeastern Germany. Scarcity of well-preserved human remains at coastal sites in these regions limits our ability to quantify the dietary importance of eelgrass resources. Ewersen et al. (2018) measured δ^13^C and δ^15^N in dog bones from several German EBC sites, but only reported individual values for Neustadt and Rosenhof, in the more saline southwest Baltic. Our analysis of their data shows that among 14 dogs with primarily marine-based diets (δ^13^C > -14‰), δ^13^C and δ^15^N are negatively correlated (p_uncorr_ 0.018, slope -0.855, r^2^ 0.446), a clear sign of the importance of eelgrass resources.

The much higher mean δ^15^N value of dogs from Baabe, on Rügen (Ewersen et al. 2018), suggests that seals may have been more important in the Baltic proper, but more detailed analysis is required. The mid-Holocene southern shoreline of the Baltic is now submerged, and it is difficult to know whether the scarcity of FHG sites with an economy focussed on marine resources east of the Darss sill (Hartz et al. 2014) reflects the lower productivity of eelgrass in the Baltic proper, or is simply a question of visibility and intensity of underwater archaeological research. In the East Baltic, seal-hunting (presumably on ice floes) was important (Lõugas 2017) but there are few coastal sites in this period, other than those relying more on lagoonal and freshwater resources (Bērziņš et al. 2022; Bērziņš et al. 2016; Piličiauskas et al. 2017; Piličiauskas et al. 2015). Stationary fishing technology was widely used inland (Piličiauskas et al. 2023), but without dense eelgrass stands, perhaps marine fish could not be harvested efficiently until the late medieval period (Orton et al. 2011).

Eelgrass productivity does not appear to explain the surprising appearance, and disappearance, of the harp seal (*Pagophilus groenlandicus*) in the mid-Holocene Baltic. Glykou et al. (2021) reported δ^13^C values for 170 prehistoric harp seals from sites around the Baltic, which are never above -15‰. Isotope data from Danish sites are not reported, but harp seals from Neustadt in the southwest Baltic have δ^13^C values as low as those in the Baltic proper, confirming that harp seal diets were based on pelagic species (e.g. herring, sprats, mature cod), and ultimately on the productivity of phytoplankton, not of eelgrass. This is unsurprising, considering the geographic distribution of harp seals in the Baltic, but confirms that isotope signatures can be used to track the use of marine resources at EBC sites (e.g. in pottery lipids).

## 4 Conclusions

Eelgrass meadows were of fundamental importance to subsistence in the Danish archipelago during the Late Mesolithic. EBC societies’ reliance on this widespread, extraordinarily productive and reliable food source may well explain their hitherto enigmatic, millennial-long coexistence with farmers in northern Germany. Eelgrass is much more productive in Danish coastal waters than in neighbouring regions of the Baltic and North Atlantic. The distinctive isotopic signature of eelgrass was transferred through the eelgrass food web, affecting isotope values in fish, marine mammals and ultimately EBC dogs and humans. This isotopic signal provides both the opportunity and the necessity to account for the contributions of eelgrass and pelagic biotopes to subsistence economies dependent on marine resources. We propose a 3-source model (eelgrass, pelagic and terrestrial resources) to improve EBC human diet reconstructions, and suggest combining this model with specific MREs in eelgrass and pelagic resources to obtain more accurate DRE-corrected dates for Ertebølle human and dog bones. We discuss a number of longstanding questions in Ertebølle research, which should be reconsidered in view of the isotopic separation of eelgrass and pelagic resources, and the productivity of eelgrass. Initial indications are that although the EBC population may have been smaller than expected, there is little to suggest that the EBC economy became unsustainable, and the end of the Ertebølle epoch may therefore have been provoked by external events rather than environmental vulnerability.

## Acknowledgments

Analyses reported here were funded by the Centre for Baltic and Scandinavian Archaeology, ZBSA research theme Man and Environment (Schleswig, Germany). AF’s contribution was supported by a professorial fellowship at the ROOTS Cluster of Excellence (Christian-Albrechts-University Kiel, Germany), funded by the Deutsche Forschungsgemeinschaft (DFG), under Germany’s Excellence Strategy (EXC 2150 – 390870439). Bones newly sampled for this study were selected and identified by Anne Birgitte Gotfredsen. We are grateful to the staff of the Leibniz Laboratory, Christian-Albrechts University, Kiel, and isolab GmbH, Schweitenkirchen, for processing and measuring the new samples. Thanks are also due to Kaare Lund-Rasmussen, Søren H. Andersen and Peter Steen Henriksen for discussing some published stable isotope values, formerly attributed to dogs.

## Supplementary information

## Appendix A. Additional information about the data sets discussed in this paper

Previously published data (δ^13^C, δ^15^N, δ^34^S and ^14^C ages) from prehistoric Danish material required to reproduce our figures and calculations are listed in Supplementary Tables 1-3. We refer readers to the original publications indicated for details of archaeological context, skeletal element, and quality control indicators. Viking-period and medieval fish δ^13^C and δ^15^N data, and supporting information, are readily accessible in the publications cited (Barrett et al. 2011; Swenson 2019).

Supplementary Table 3 includes δ^13^C, δ^15^N and ^14^C ages for human skeletal remains from Denmark dating to the Mesolithic, including individuals of the transitional Mesolithic-Neolithic period with a 100% Western Hunter-Gatherer genetic profile (Allentoft et al. 2024). In four cases, more than one sample of the same individual has been analysed, and we arbitrarily selected only one of these samples to avoid introducing bias to the overall data set. When different samples of the same individual give δ^13^C, δ^15^N and/or ^14^C ages differences that cannot be attributed to contamination or laboratory errors, they may reveal lifetime changes in diet, as different skeletal elements (dentine, petrous bone, cortical bone, trabecular bone) are remodelled at different rates (van der Sluis et al. 2015). Intra-skeletal differences in δ^13^C, δ^15^N and ^14^C age potentially represent opportunities to refine the 3-source DRE correction model presented in this paper (3.3.3, Meadows et al. (2024)).

The material presented here (Table 1) was obtained from various projects over several years. General information about the sites mentioned in Table 1 is already published (Asnæs Havnemark: Price et al. (2018); Bodal K: Fischer and Gotfredsen (2006); Fischer et al. (2007a); Olsbjerg: Liversage (1973); Østenkær: Lysdahl (1985); Rødhals: Fischer et al. (2021)). Ten of the twelve samples are from sites located at the contemporaneous coastline (Asnæs Havnemark, Olsbjerg, Rødhals).

Zoological experts of the Quaternary Zoological Department of Denmark’s Nature Historical Museum in Copenhagen were responsible for the selection of bones to be sampled, ensuring that no animal was unwittingly sampled twice. This selection was based on anatomical part, symmetry and size/biological age. For this intricate work, we are much indebted to Knud Rosenlund (deceased 2023), Inge Bødker Enghoff and Anne Birgitte Gotfredsen.

## Appendix B. Human diet reconstruction and DRE-correction models

In order to visualise the importance of each biotope to individual human diets (Fig. 10) and estimate each biotope’s contribution to human ^14^C ages, we use the Bayesian statistical package FRUITS 2 (Fernandes et al. 2014), which iteratively compiles potential dietary mixtures that could produce the measured isotope values in each individual, based on representative isotope values for food sources and diet-collagen fractionation. The output (Supplementary Table 4) includes posterior probability distributions for the overall intake of each food source by each individual, and the contribution of each food source to each isotope value. These are identical in an unrouted (concentration-independent) FRUITS model, i.e. if we assume that macronutrient concentrations were similar in each food source. As each food source’s percentage contribution to collagen ^13^C and ^14^C contents must be identical, we can use FRUITS output for dietary reservoir effect (DRE) correction, which facilitates the propagation of uncertainties.

Our FRUITS model parameters are based on empirical data (Supplementary Table 1; (Fischer et al. 2007b; Maring and Riede 2019)), with appropriate isotopic spacing from faunal collagen to flesh protein (Fernandes et al. 2015; Fischer et al. 2007a) to obtain representative δ^13^C and δ^15^N values for dietary protein. For the terrestrial biotope, isoscapes are relatively well understood (with the exception of δ^34^S, which we are not currently using in diet reconstruction), and δ^13^C and δ^15^N appear to vary little between species. Simply averaging faunal isotope data is valid, as wild woodland herbivores should not have access to pelagic or eelgrass food webs (c.f. Maring and Riede 2019). The terrestrial dietary protein values applied here would need to be badly wrong to significantly affect diet reconstructions, considering the overwhelmingly marine diets of most EBC individuals, and the isotopic distances between the terrestrial biotope and both marine biotopes.

Dietary protein values for 100% pelagic or eelgrass biotope food sources are necessarily notional, because most marine faunal isotope values represent a continuum between phytoplankton and eelgrass primary productivity. For the pelagic biotope, archaeological faunal data include fish taxa whose δ^13^C and δ^15^N values are not obviously influenced by the eelgrass biotope, such as mackerel, and marine mammals which appear to have depended overwhelmingly on pelagic fish. These taxa span much wider δ^13^C and δ^15^N ranges than terrestrial herbivores, partly due to trophic-level fractionation, and perhaps metabolic factors in seals, so the main barrier to defining average dietary protein δ^13^C and δ^15^N values is rationalising the weight given to each taxon. Marine mammal bones may be more visible and less prone to diagenesis, but we assume that herrings were consumed more often, even if they are entirely absent from the published prehistoric isotope data.

Even fish taxa that are predominantly associated with the eelgrass biotope (eels and flatfish) show negative δ^13^C-δ^15^N correlations, which imply a varying degree of dependence on the pelagic biotope. Thus simply averaging their isotope values cannot provide valid isotope values for dietary protein derived only from eelgrass primary production. The values applied here are therefore subjective estimates, informed by δ^13^C and δ^15^N values in ‘marine’ dogs (Fig. 5). Several dogs and one otter have δ^13^C values of -9 to -8‰, which may represent ∼100% eelgrass-biotope diets. Dogs appear to have had more specialised or monotonous diets than humans, partly because humans are much longer-lived, and most human samples represent average diets over a number of years.

Table B1 shows the isotope values and macronutrient concentrations we regard as realistic. Plant and mollusc values for the EBC are not measured directly, but can be inferred from collagen isotope values, and modern isotope spacing and nutrient profiles.

**Table B1.** Assumed δ^13^C and δ^15^N values and macronutrient concentrations for food sources.

| original values in food sources | collagen |  | edible protein |  | macronutrients |  |
| --- | --- | --- | --- | --- | --- | --- |
| food | $\delta^{13}\text{C}$ (‰) | $\delta^{15}\text{N}$ (‰) | $\delta^{13}\text{C}$ (‰) | $\delta^{15}\text{N}$ (‰) | protein | energy |
| terrestrial herbivore | -22.0 | 5.5 | -24.5 | 7.0 | 70% | 30% |
| plant |  |  | -26.5 | 2.5 | 10% | 90% |
| seal, cetacean | -14.5 | 14.0 | -17.0 | 15.5 | 40% | 60% |
| pelagic fish | -15.5 | 11.0 | -18.0 | 12.5 | 60% | 40% |
| eelgrass fish | -10.0 | 7.5 | -12.5 | 9.0 | 40% | 60% |
| marine mollusc (flesh) |  |  | -13.5 | 5.5 | 60% | 40% |

For the FRUITS model, these values have been simplified by taking weighted averages of values from the same biotope, resulting in only 3 food sources, ‘terrestrial’, ‘pelagic’ and ‘eelgrass’ (Table B2). We apply smaller uncertainties to terrestrial isotope values, as terrestrial herbivore isotope data are less scattered than faunal data from either of the marine biotopes. We assume that weighted average macronutrient concentrations are similar enough in foods from each biotope that we can ignore any effect on δ^13^C of lipids and carbohydrates by using an unrouted FRUITS model (Fernandes 2016). We apply trophic enrichment factors (diet-collagen) of +4.5±0.5‰ and +5.0±0.5‰ for δ^13^C and δ^15^N respectively (Fernandes et al. 2015).

**Table B2.**
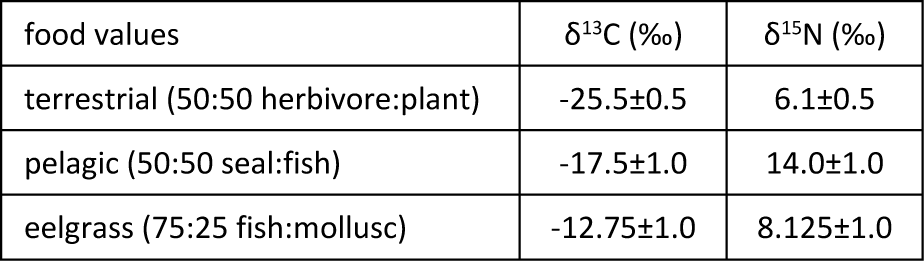
Weighted average δ^13^C and δ^15^N values in foods from each biotope.

Figure B1 plots Danish Mesolithic human δ^13^C and δ^15^N values (Supplementary Table 3, Fig. 8) against an expected isotope range for human consumers of known and monotonous diets, obtained using the parameter values applied in the FRUITS model. The fact that (nearly) all individuals fall within this range is a good indication that these parameter values are realistic. Any model which merges pelagic and eelgrass resources into a single marine food group (i.e. with fixed proportions of pelagic and eelgrass species) produces a much narrower expected isotopic range, excluding most of the individuals analysed (Fig B1). A handful of cases have lower δ^13^C and/or higher δ^15^N values than expected if they only consumed foods from the terrestrial, pelagic and eelgrass biotopes. In these cases, freshwater fish (typically having lower δ^13^C and/or higher δ^15^N than terrestrial or marine species, e.g. Robson et al. (2016)) was probably a major component of diet. Individuals falling within our 3-source model’s expected range may have consumed some freshwater fish, but their isotope values are also consistent with negligible freshwater fish consumption. Any freshwater fish intake would displace more of the terrestrial food intake indicated in our 3-source model output, than the pelagic or eelgrass-biotope food intake.

**Figure B1.**
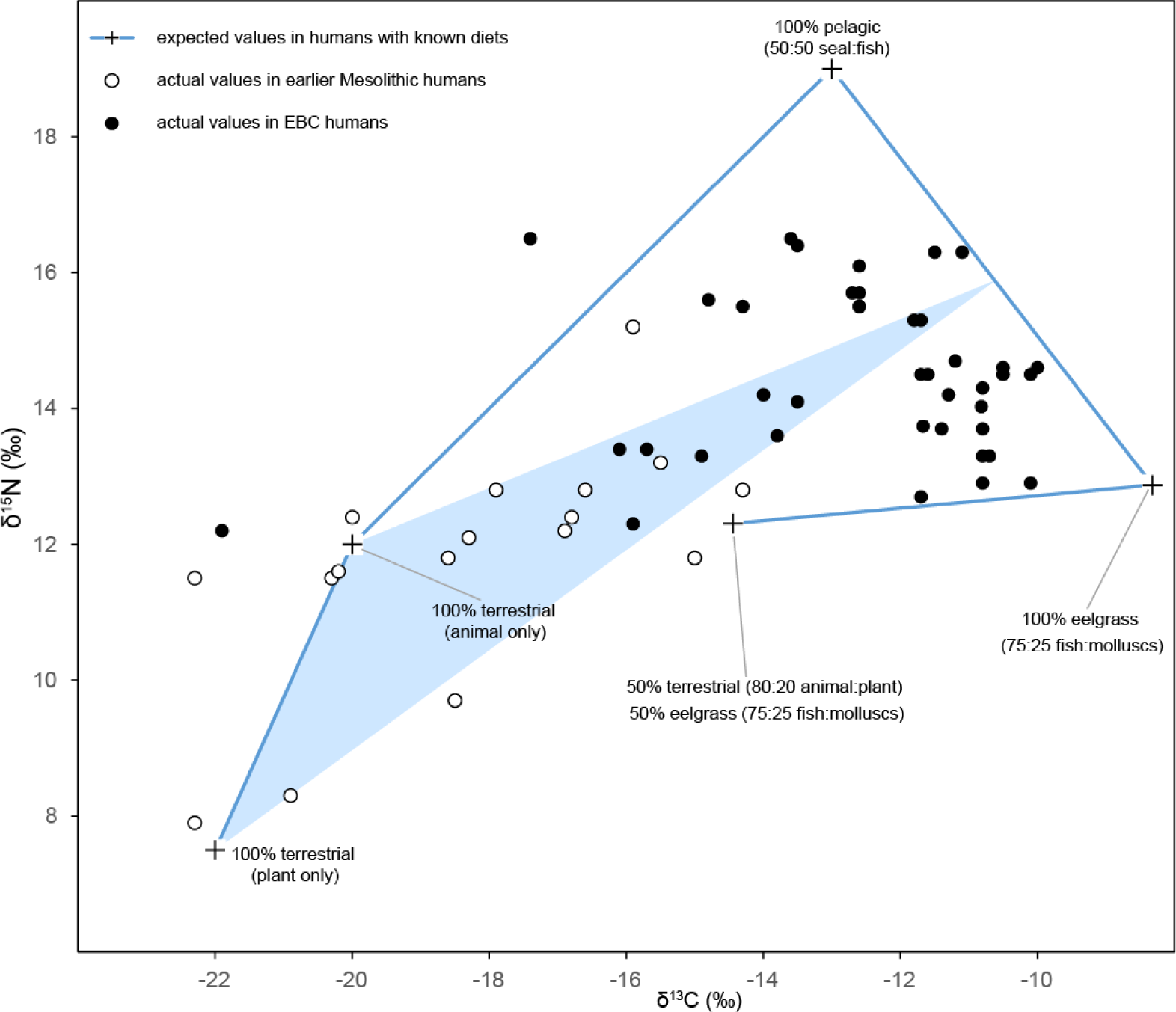
Danish Mesolithic human collagen δ^13^C and δ^15^N values (Allentoft et al. 2024; Allentoft et al. 2022; Fischer et al. 2007a; Price et al. 2007) plotted against a potential range obtained using the parameter values employed in the FRUITS model, and hypothetical individuals (+) with monotonous diets. The shaded area indicates the expected range of δ^13^C and δ^15^N values if diets are mixtures of terrestrial plants and animals, and a single marine food group composed of equal amounts of pelagic and eelgrass resources.

To calibrate ^14^C ages subject to two DREs, we define separate calibration curves for pelagic and eelgrass food sources (Marine20 (Heaton et al. 2020) with a Delta_R of 0±50y and -264±61y respectively; see Appendix C). We then use the OxCal Mix_Curves function (Bronk Ramsey 2009) twice for each date. This is illustrated by an example:

~~~
Mix_Curves("AAR-8552 marine", "pelagic", "terrestrial", 24.2, 7.8);
Mix_Curves("AAR-8552 coastal","AAR-8552 marine", "eelgrass", 85.1, 9.5);
R_Date("AAR-8552 Rødhals, I", 5360, 50);
~~~

The first line of code defines the contribution of the ‘terrestrial’ calibration curve (IntCal20, Reimer et al. (2020)), based on the FRUITS model’s mean estimate of terrestrial protein intake for this individual, 24.2±7.8%; the marine component is calibrated using the ‘pelagic’ calibration curve. The second line recalibrates the marine component with a 85.1±9.5% contribution from the ‘eelgrass’ calibration curve. This percentage is not simply the FRUITS mean estimate of protein intake from the eelgrass biotope (as a percentage of the total diet), but its estimate of the proportion of marine protein derived from the eelgrass biotope, which is calculated (‘eelgrass % of marine’, Supplementary Table 4) from the mean estimates of protein intake from all 3 sources. We estimate the uncertainty in ‘eelgrass % of marine’ by scaling the average uncertainty of each marine food source contribution to overall diet (calculated as 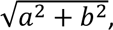 where *a* = sd%_pelagic_ and *b* = sd%_eelgrass_) to the combined marine contribution to overall protein intake (1-mean%_terrestrial_).

In individuals with a more terrestrial diet, the absence of a freshwater food group in the FRUITS model may lead to under-estimation of the amount of marine protein consumed, because freshwater fish δ^13^C values were probably lower than the terrestrial herbivore δ^13^C values. More importantly, any freshwater fish intake would add another DRE, which is difficult to quantify, due to the scarcity of reference stable isotope and ^14^C data from freshwater biotopes in prehistoric Denmark. Freshwater reservoir effects may exceed those in marine biotopes (Maring et al. 2024) and DRE corrections given by our 3-source model for the Tingbjerggård Vest EBC individual, and earlier Mesolithic individuals (Supplementary Table 4), are therefore probably under-estimates. As this problem barely affects most EBC individuals, we have not attempted to resolve it here.

## Appendix C. Uncertainty and variability in applicable MREs

As discussed in the main text (3.3.3), there is little empirical evidence to constrain the MRE in pelagic species from mid-Holocene Denmark. The small number of post-medieval cetaceans dated (Mangerud et al. 2006; Olsson 1980) may not be relevant to the Danish situation and in the British Isles, mid-Holocene MREs appear to have been higher than in the medieval period (Ascough et al. 2009). For illustrative purposes, therefore, we have applied the global surface ocean calibration model, Marine20 (Heaton et al. 2020), with a small additional uncertainty (ΔR of 0±50y) to calibrate the pelagic component of human diets.

To calibrate the eelgrass component, we calculated ΔR values relative to Marine20 using published ^14^C data from marine shells, primarily oysters, at four coastal sites: Asnæs Havnemark (Price et al. 2018), Hjarnø (Astrup et al. 2021), Horsens Fjord and Tempelkrog (Olsen et al. 2017; Olsen et al. 2009), on the basis that these shells were formed from the same DIC available to eelgrass for photosynthesis. We used Bayesian chronological models (Bronk Ramsey 2009) to constrain the range of ΔR values required to synchronise these shells with terrestrial samples from the same sites.

The OxCal CQL code given below exactly specifies the ^14^C data and chronological interpretations incorporated in each site model. For simplicity, these site models are included as independent sequences in a single OxCal model. The first part of the OxCal model generates likelihoods for the dates of shell samples by calibrating their ^14^C ages using Marine20 with a vague likelihood for the associated ΔR, e.g.:

~~~
Delta_R("609.5cm Ostrea edulis", N(-300,100));
R_Date("609.5cm Ostrea edulis AAR-9688", 2301, 46);
~~~

The likelihood for each shell date is then cross-referenced at the appropriate point in the corresponding site sequence. We are primarily interested in the posterior distributions of ΔR likelihoods for Ertebølle-epoch shells. At two sites, Horsens Fjord and Tempelkrog (Olsen et al. 2017; Olsen et al. 2009), the dated shells are from stratified sequences spanning much of the Holocene, and we apply independent ΔR likelihoods for the earliest and latest shells, in case the eelgrass biotope MRE changed over time. As these were natural sedimentary sequences, we used the OxCal P_Sequence function to model the site chronologies (Bronk Ramsey 2009). We use published stratigraphic evidence to constrain the dates of samples from the archaeological sequences at Asnæs Havnemark (Price et al. 2018) and Hjarnø (Astrup et al. 2021). Our model code also reproduces (Fischer and Olsen 2021)’s model to estimate ΔR in closely dated Neolithic oysters from Nekselø. This estimate (-234±61y) appears to be fairly representative of the Ertebølle-epoch shell MREs at the other four sites. We have not included a model to recalculate ΔR at Syltholm, because the data given by Philippsen (2018) consist mainly of bulk sediment ^14^C ages, but the Nekselø ΔR would also fit here (under reasonable assumptions about the sources of organic carbon in bulk sediment). We have therefore adopted this ΔR estimate for our eelgrass biotope calibration curve.

## Footnotes

1 If freshwater δ^34^S is 6‰ and freshwater contains as much sulfate as oceanic saltwater, δ^34^S will be ∼15‰ in a 1:1 mix (i.e. salinity of 17-18‰). For δ^34^S to be <10‰, either freshwater δ^34^S must be much lower, or salinity must be much lower, or freshwater sulfate concentration must be much higher. As seals in the East Baltic (salinity <10‰) have δ^34^S of ∼16‰ (Aguraiuja-Lätti et al. 2022), however, the sulfate concentration in freshwater entering the Baltic is apparently much lower than in oceanic saltwater.

2 We disregard δ^34^S data from dogs and several other mammal species at Syltholm, as even among δ^34^S data accepted by Gron et al. (2024), δ^34^S is significantly correlated with %S, C/S and N/S (for all taxa or dogs only, puncorr 0.002 to 0.003), suggesting contamination with low-δ^34^S sulfur from the burial environment.

3 Including an individual at inland Tingbjerggård Vest, whose diet was based on terrestrial and/or freshwater foods (δ^13^C -21.9‰, δ^15^N 12.2‰), but not an undated human bone from Gudumlund (δ^13^C -10.6‰, δ^15^N 13.1‰ (van der Sluis et al., 2019)) which may have been attributed to the EBC due to its isotope values.

